# Adaptive protein coevolution preserves telomere integrity

**DOI:** 10.1101/2024.11.11.623029

**Authors:** Sung-Ya Lin, Hannah Futeran, Mia T. Levine

## Abstract

Many essential conserved functions depend, paradoxically, on proteins that evolve rapidly under positive selection. How such adaptively evolving proteins promote biological innovation while preserving conserved, essential functions remains unclear. Here, we experimentally test the hypothesis that adaptive protein-protein coevolution within an essential multi-protein complex mitigates the deleterious incidental byproducts of innovation under pressure from selfish genetic elements. We swapped a single, adaptively evolving subunit of a telomere protection complex from *Drosophila yakuba* into its close relative, *D. melanogaster*. The heterologous subunit uncovered a catastrophic interspecies incompatibility that caused lethal telomere fusions. Restoring six adaptively evolving sites on the protein-protein interaction surface, or introducing the *D. yakuba* interaction partner, rescued telomere integrity and viability. Our *in vivo*, evolution-guided manipulations illuminate how adaptive protein-protein coevolution preserves essential functions threatened by an evolutionary pressure to innovate.

## Main Text

Strictly conserved proteins support strictly conserved functions. For example, the ancient and conserved heat shock protein, Hsp70, determines a similarly conserved stress response mechanism across eukaryotes (*1*). Likewise, rapidly evolving proteins support rapidly evolving functions. The rapidly evolving immune factor, Protein kinase R, determines species-specific responses to poxvirus infection in primates (*2*). Surprisingly, many rapidly evolving proteins also support strictly conserved, essential functions. Rapid evolution under positive selection is pervasive across essential centromere proteins that support chromosome segregation (*3, 4*), DNA repair proteins that maintain genome integrity (*5, 6*), and nuclear pore proteins that regulate trafficking across the nuclear membrane (*7, 8*). Myriad computational and *in vitro* experimental studies offer a compelling resolution to this paradoxical observation: amino acid changes in one protein trigger compensatory changes in interacting proteins, RNA, or DNA (*9-21*). This coevolution may mitigate the deleterious incidental byproducts of evolutionary innovation in response to a potent evolutionary pressure. However, *in vivo* experimental investigations of compensatory evolution to preserve essential functions are limited to deeply divergent species that preclude inferences of adaptive evolution (*22, 23*). Consequently, whether and how adaptive coevolution preserves essential functions remains unclear (*24*), and yet disrupted coevolution may underlie many lethal F_1_ hybrid incompatibilities (*25-29*).

The *Drosophila* telomere offers an ideal model to investigate evolutionary innovation constrained by an essential function. A six-subunit “end-protection complex” binds to telomeric DNA in a sequence-independent manner and prevents inappropriate recognition of the terminal DNA as a double-strand break (*30, 31*). Depletion of any one subunit results in lethal end-to-end chromosome fusions (*32-38*). This vital protection of chromosome ends is established immediately following fertilization. Sperm-deposited paternal chromosomes initially lack the end-protection complex (*39-41*). Maternally provisioned subunits in the embryo assemble into a complex at paternal telomeres (*39*), ensuring paternal genome integrity and faithful chromosome segregation. Once established, the end-protection complex is epigenetically propagated through development. Despite the essentiality and universal conservation of linear chromosome end-protection, two of the six subunits of the *Drosophila* end-protection complex evolve adaptively (*5, 42*) (Fig. 1A, Table S1 and S2). We previously showed that one such protein, HOAP (HP1/ORC-Associated Protein), recurrently evolves to restrict the deleterious proliferation of telomeric retrotransposons (*42*). Evidence of ongoing antagonistic coevolution with retrotransposons indicates that this essential, multi-protein complex is under pressure to innovate. Intriguingly, the only other adaptively evolving subunit in this complex is a protein called HipHop (HP1-HOAP-interacting protein, Fig. 1A, Table S1 and S2). HipHop and HOAP physically interact and are interdependent for stability and telomere localization (*33*). These data raise the possibility that adaptive coevolution between these two essential telomere-binding proteins is required to preserve complex integrity and chromosome end-protection. Under this model of adaptive coevolution, a species-specific version of HipHop requires a species-specific version of HOAP to support telomere integrity. HipHop from one species should be incompatible with HOAP from even a closely related species.

**Figure 1.**
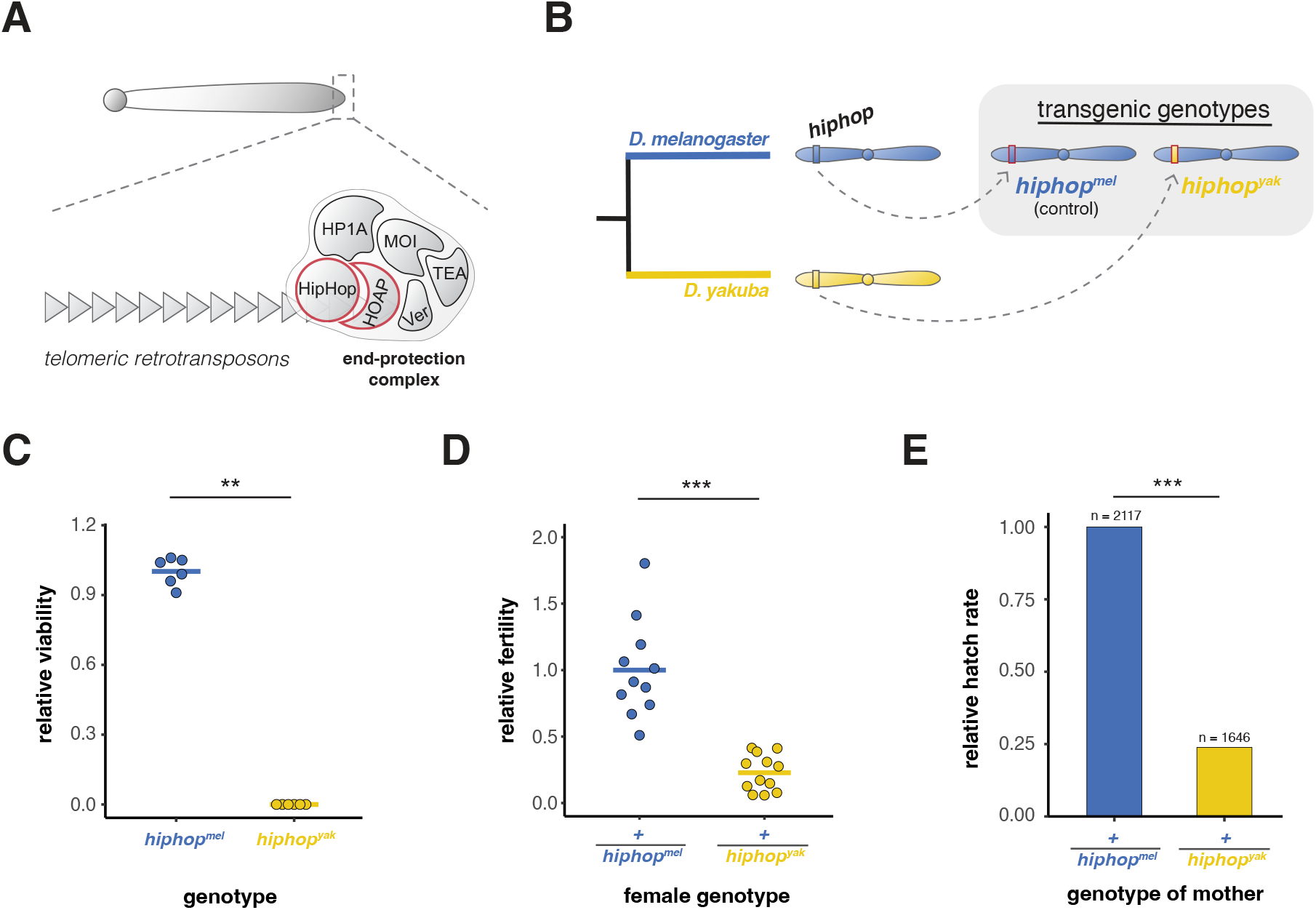
Adaptive protein evolution of HipHop results in a catastrophic cross-species incompatibility. **(A)** A schematic of the *Drosophila* telomere (*30, 31*). Red outline indicates adaptively evolving subunits of the end-protection complex. (**B**) Schema of the CRISPR/Cas9-mediated gene swap approach. We replaced the native *hiphop* coding sequence with either the *D. melanogaster hiphop* (“*hiphop^mel^*”) or the *D. yakuba hiphop* (“*hiphop^yak^*”) intron-less coding sequence, tagged with EGFP (rectangle outlined in red). (**C**) Relative viability of *hiphop^mel^* and *hiphop^yak^* (Mann-Whitney U test: \*\**p*-value < 0.01). (**D**) The relative female fertility of *+/hiphopmel* and *+/hiphopyak* females crossed to wild-type males (MWU: \*\*\**p*-value < 0.001). (**E**) Relative hatch rate of embryos from *+/hiphopmel* and *+/hiphopyak* females. (Fisher’s Exact Test: \*\*\**p*-value < 0.001). “*+*” = *TM6* balancer chromosome.

To experimentally test for an inter-species incompatibility, we replaced the endogenous *D. melanogaster hiphop* coding sequence with a highly diverged, EGFP-tagged version from its close relative, *D. yakuba* (*hiphop*^*yak*^) (Fig. 1B). In parallel, we engineered a control genotype with an EGFP-tagged version of the *D. melanogaster* coding sequence (*hiphop*^*mel*^, Fig. 1B). If *hiphop*^*yak*^ is incompatible with the *D. melanogaster* genetic background, the *D. yakuba* version of HipHop is expected to reduce viability and/or fertility. We discovered that the wildtype allele of *hiphop* from *D. yakuba* is homozygous lethal in *D. melanogaster* (Fig. 1C). Homozygous *hiphop*^*yak*^ individuals developed to the third instar larval stage, but most failed to pupate (Fig. S1A). Moreover, the *+/hiphop*^*yak*^ heterozygous individuals were viable; however, females were severely subfertile (Fig. 1D, but not males, Fig. S1B). Heterozygous female subfertility was unexpected because *hiphop* heterozygous null females suffer no such fertility loss (Fig. S1C).

Appreciating that maternally provisioned HipHop is required to establish telomeres in the early embryo (*39*), we wondered if these *+/hiphop*^*yak*^ females’ embryos, rather than their egg production, accounted for the subfertility. Consistent with this prediction, we observed no reduction in egg number (Fig. S1D) but a severely reduced embryo hatch rate in the progeny of *+/hiphop*^*yak*^ heterozygous females (Fig. 1E). These data suggest that *hiphop*^*yak*^ is both homozygous lethal (Fig. 1C) and dominant maternal effect lethal – maternally provisioned HipHop^yak^ disrupts embryogenesis despite maternal provisioning of endogenous (“*+*”) HipHop^mel^ as well.

To determine if loss of chromosome end-protection underlies the observed lethal inter-species incompatibility, we assayed telomere-telomere fusion rates in the homozygous *hiphop*^*yak*^ larva, the developmental stage immediately before lethality. At this late larval stage, the percentage of telomere fusions in *hiphop*^*yak*^ was significantly elevated relative to the *hiphop*^*mel*^ control (Fig. 2A). These data indicate that chromosome end-protection fails in *hiphop*^*yak*^ homozygotes.

**Figure 2.**
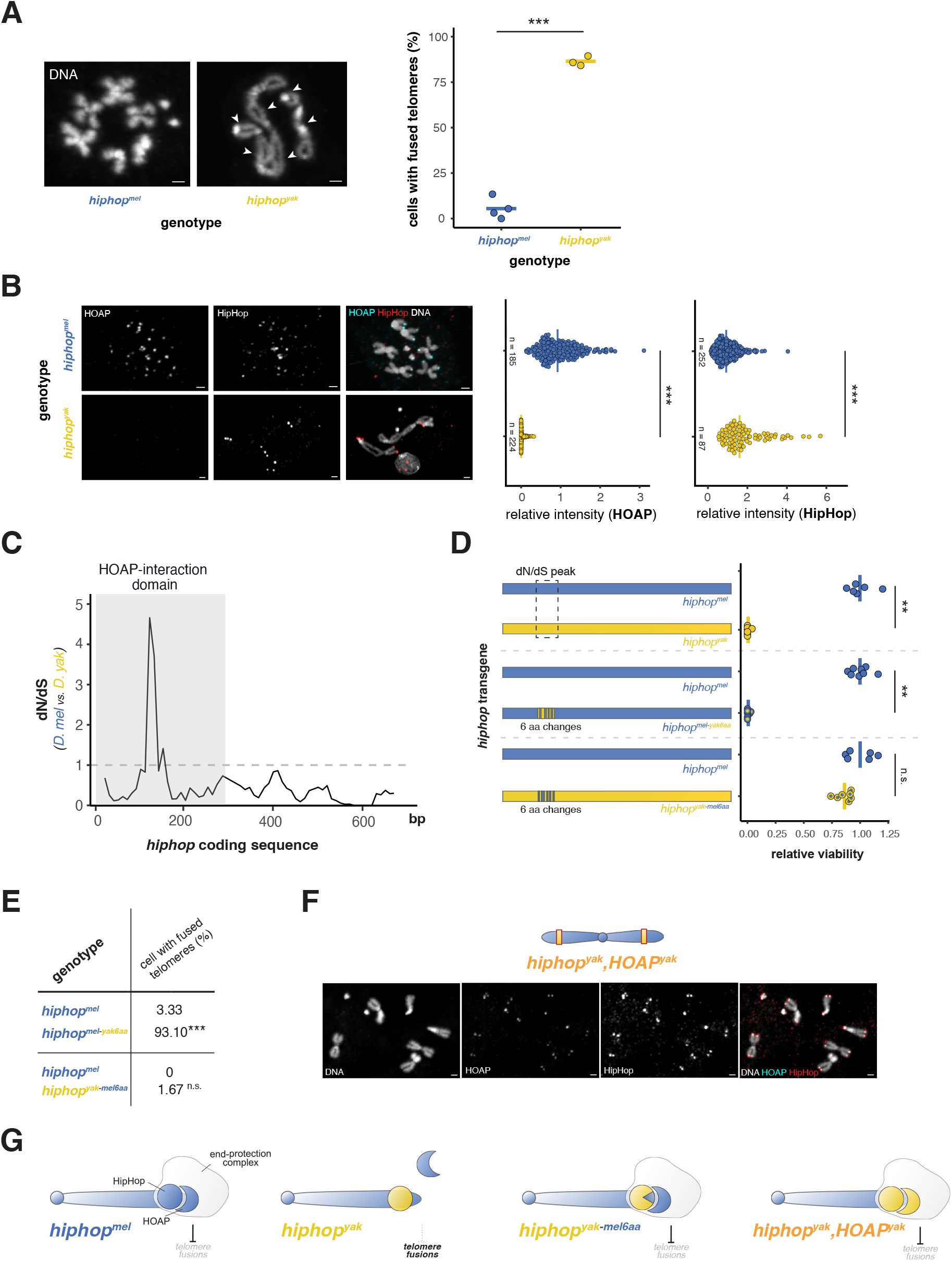
Adaptive evolution of the HipHop-HOAP interaction surface preserves telomere protein recruitment and end-protection. (**A**) Images of mitotic chromosomes from larval brains (left, arrowheads indicate fusions) and the percent mitotic cells with telomere fusions observed in *hiphop^mel^* and *hiphop^yak^* homozygotes (right). Each data point represents >100 mitotic cells scored from one individual. Fisher’s Exact Test, \*\*\**p*-value < 0.001. (**B**) Mitotic chromosomes from larval brains stained for HOAP (anti-FLAG) and HipHop (anti-EGFP) (left). Quantification of the HOAP-FLAG and EGFP-HipHop intensity at *hiphop^mel^* and *hiphop^yak^* homozygote telomeres, normalized by mean intensity of *hiphop^mel^* (right). MWU test, \*\*\**p*-value < 0.001. (**C**) The estimated rate of nonsynonymous substitutions relative to synonymous substitutions (dN/dS) across the *hiphop* coding sequences between *D. melanogaster* and *D. yakuba* (window size = 50, step size = 10). (**D**) Relative viability of *D. melanogaster*-*D. yakuba* chimeric *hiphop* homozygous adults. The dashed box indicates the *hiphop* coding region with elevated dN/dS (see Figure 2C). All data points normalized by the mean viability of each *hiphop^mel^* control strain into which the respective transgene was introgressed. MWU test, \*\**p*-value < 0.01 (**E**) Percent mitotic cells with telomere fusions observed in *D. melanogaster*-*D. yakuba* chimeric *hiphop* homozygotes. n ^3^ 58. Fisher’s Exact Test, \*\*\**p*-value < 0.001. (**F**) Schema of chromosome encoding *hiphop^yak^* recombined with a previously constructed swap (*42*) of 3xFLAG-tagged, *D. yakuba cav/HOAP* coding sequence into *D. melanogaster* (above). Mitotic chromosomes from larval brains observed in *hiphop^yak^,HOAPyak* homozygotes stained for HOAPyak (anti-FLAG) and HipHopyak (anti-EGFP, below). (**G**) Model of telomere end-protection complex assembly and suppression of telomere fusions in the presence of HipHopmel, HipHopyak, HipHopyak-mel6aa, and HipHopyak,HOAPyak in *D. melanogaster*. Scale bars, 1 μm.

In *D. melanogaster*, the localization of both HipHop and HOAP to telomeres is required for chromosome end-protection (*32, 33*), and the two proteins are interdependent for stability (*33*). We hypothesized that failure of HipHop and HOAP to localize to telomeres causes loss of end-protection in *hiphop*^*yak*^ homozygotes. In the developmental stage where we observed elevated telomere fusions (Fig. 2A), we detected no *D. melanogaster* HOAP at the telomeres of *hiphop*^*yak*^ homozygotes (Fig. 2B, left). Given that HipHop depends on HOAP for stability in *D. melanogaster* (*33*), we also predicted that HipHop^yak^ would fail to localize to telomeres. Contrary to our prediction, HipHop^yak^ localized robustly despite the absence of HOAP (Fig. 2B, left). In fact, we detected significantly more HipHop^yak^ than HipHop^mel^ at the *D. melanogaster* telomeres (Fig. 2B, right). Robust, HOAP-independent localization of HipHop^yak^ suggests that *D. yakuba* employs a distinct mechanism of localization.

The failure of *D. melanogaster* HOAP to localize to the telomeres of *hiphop*^*yak*^ flies supports the possibility that HipHop evolves adaptively to preserve its interaction with HOAP and protect telomeres from fusions. Under this model, adaptive sequence evolution may have shaped specifically the interaction surface of HipHop and HOAP. To investigate this possibility, we estimated the ratio of nonsynonymous to synonymous divergence (dN/dS) along the *hiphop* coding sequence between *D. melanogaster* and *D. yakuba*. Consistent with a history of positive selection shaping this protein-protein interaction, we observed a single peak of dN/dS within the HOAP-interaction domain of HipHop (Fig. 2C). Only six HipHop residues underlie the narrow peak of positive selection (Fig. 2D and S2A).

To test the possibility that the evolution of at most six residues is necessary and sufficient for viability, we replaced the endogenous *D. melanogaster* HipHop with *D. melanogaster*-*D. yakuba* chimeric HipHop proteins (Fig. 2D). We first engineered a fly with the six *D. yakuba* residues in an otherwise *D. melanogaster* version of the protein. These flies displayed elevated telomere fusions (Fig. 2E and S2B) and were homozygous lethal (Fig. 2D), largely around the larva-to-pupa transition. We also engineered the reciprocal *D. melanogaster* fly with the six *D. melanogaster* residues in an otherwise *D. yakuba* version of the protein. These flies displayed well-resolved telomeres (Fig. 2E and S2B) and were homozygous viable (Fig. 2D). Together, these data suggest that the evolution of at most six residues on the HipHop-HOAP interaction surface is both necessary and sufficient for end-protection and adult viability.

Under a model of HipHop-HOAP coevolution, swapping these six amino acids from *D. melanogaster* into the *D. yakuba* HipHop is one of two possible approaches to reverse *hiphop*^*yak*^*-* mediated incompatibility. The *D. yakuba* version of HipHop’s interaction partner, HOAP, should similarly restore telomere protection in *hiphop*^*yak*^. Consistent with this prediction, introducing *HOAP*^*yak*^ into the *hiphop*^*yak*^ genotype restored HOAP localization to telomeres (Fig. 2F), restored telomere end-protection (Fig. 2F), and restored viability (Fig. S3A). This robust rescue indicates that HOAP alone is responsible for the cross-species incompatibility triggered by HipHop^yak^ (Fig. 2G), and offers the first empirical evidence of adaptive protein-protein coevolution preserving an essential function.

Our aim to experimentally test, *in vivo*, a classic model of protein-protein coevolution yielded discoveries only partially consistent with the model’s predictions. The classic model of coevolution predicts that replacing one protein with a heterologous version from a closely related species leads to loss of protein-protein interaction and, consequently, loss of function (*24, 43*). Indeed, the recessive loss-of-function phenotype of *hiphop*^*yak*^ mirrored the null mutant, which, in this case, is recessive lethality associated with loss of end-protection (*33, 39, 40*). However, unlike the coevolution model predictions, we also observed that *hiphop*^*yak*^ can act dominantly (Fig. 1D and 1E). Heterozygous *+/hiphop*^*yak*^ females are dominant maternal effect lethal – embryos from these mothers were inviable. Notably, HipHop not only maintains telomeres through development (*33*) but also epigenetically establishes telomeres on the sperm-deposited paternal chromosomes (*39, 40*) in the embryo. During spermiogenesis, the canonical end-protection complex is replaced by at least two sperm-specific telomere-binding proteins (*39-41*). In the zygote, these paternal chromosome-specific telomere proteins are replaced by maternal HipHop and other maternally provisioned members of the complex (*39*). We wondered if maternal failure to establish paternal telomere end-protection in the early embryo might account for the observed dominant maternal effect lethality of *+/hiphop*^*yak*^ heterozygous females (Fig. 1E).

To investigate the basis of this dominant lethality, we characterized early embryonic development. We observed a high percentage of embryos from *+/hiphop*^*yak*^ heterozygous mothers that failed to reach the developmental stage when the zygotic genome initiates transcription (zygotic genome activation, or ZGA) (*44*) (Fig. 3A). This finding is consistent with our inference of maternal effect lethality: maternal provisioning, rather than the zygotic factors, caused embryonic lethality. To determine the stage of this early embryonic arrest, we visualized early embryos from *+/hiphop*^*mel*^ and *+/hiphop*^*yak*^ mothers and assessed the distribution of early embryonic stages. We observed strikingly different frequency distributions of embryonic development from *+/hiphop*^*mel*^ and *+/hiphop*^*yak*^ mothers. Embryos from *+/hiphop*^*yak*^ mothers were especially enriched at nuclear cycle 1 (Fig. 3B, FET, *p* <0.001), consistent with very early embryonic arrest.

**Figure 3.**
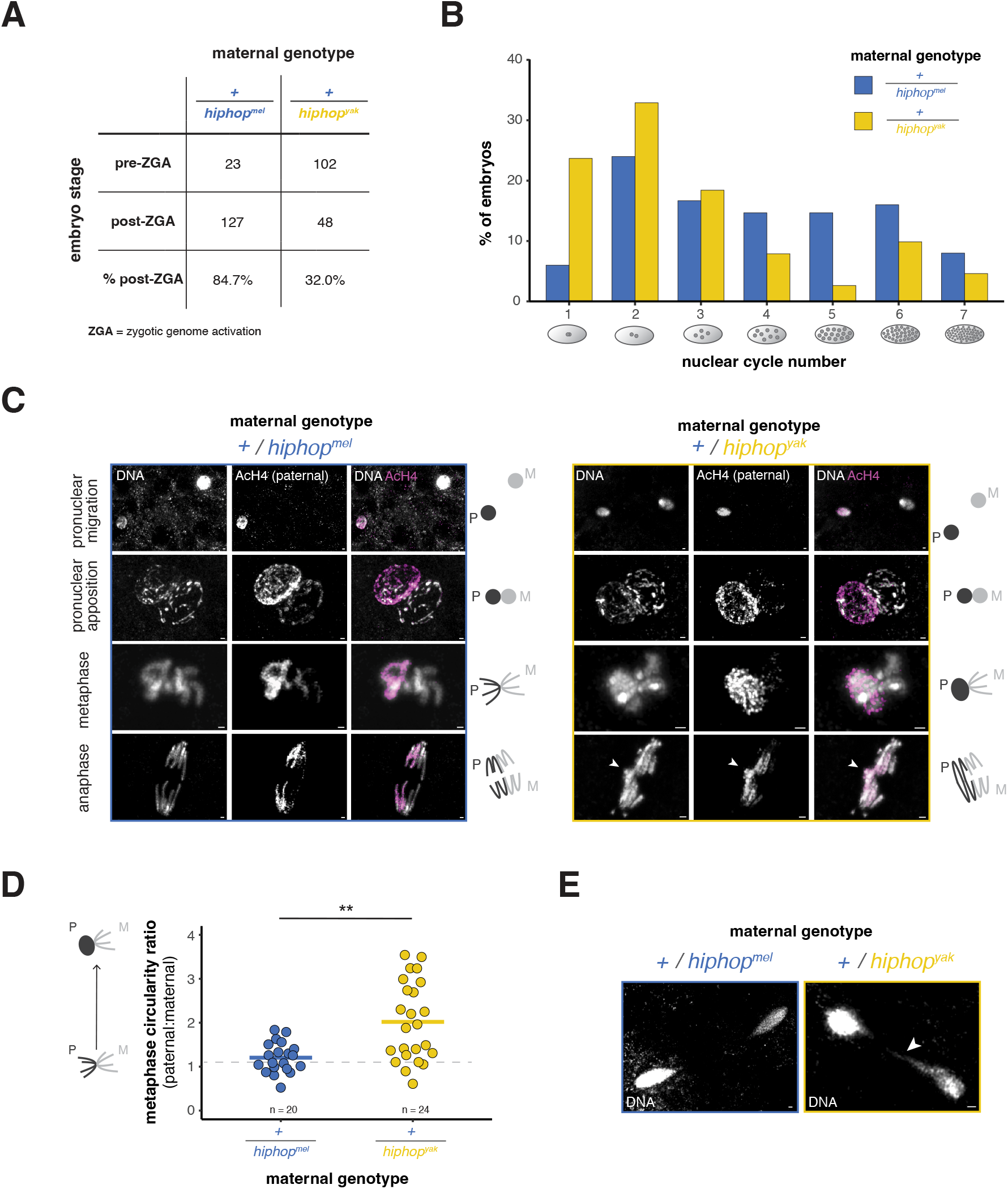
HipHopyak interferes with the establishment of paternal telomere protection in the zygote. (**A**) Percent of embryos from +/*hiphop^mel^* and +/*hiphop^yak^* females that develop beyond nuclear cycle 14, which marks the major wave of zygotic genome activation (ZGA). n = 150. Fisher’s Exact Test, *p*-value < 0.001. (**B**) Percent of embryos at nuclear cycle numbers 1-7 from a 1-hour collection from +/*hiphop^mel^* and +/*hiphop^yak^* females. n ^3^ 150. Fisher’s Exact test, *p*-value < 0.001. (**C**) Zygotic chromosomes before or during the first mitosis (with schematics on the right) from +/*hiphop^mel^* and +/*hiphop^yak^* females. Acetylated H4 (AcH4) marks the paternal chromatin. The arrowhead indicates a paternal chromatin bridge. P = paternal, M = maternal. (**D**) Circularity ratio of paternal and maternal chromatin at metaphase in embryos from +/*hiphop^mel^* and +/*hiphop^yak^* females. MWU test, \*\**p*-value < 0.01. P = paternal, M = maternal. (**E**) Zygotic chromosomes at the first telophase from +/*hiphop^mel^* and +/*hiphop^yak^* females. The arrowhead indicates a chromatin bridge.“*+*” = *TM6* balancer chromosome for all panels. Scale bars, 1 μm.

To test the hypothesis that developmental arrest arises from a failure to establish end-protection on paternal chromosomes, we stained nuclear cycle 1 embryos with a marker of paternal chromatin (acetylated histone H4, AcH4). The earliest detectable defect in embryos from *+/hiphop*^*yak*^ mothers occurred at metaphase. Unlike the maternal chromosomes, the paternal chromosomes failed to condense at metaphase (Fig. 3C and 3D). At anaphase, we detected a paternal chromatin bridge (Fig. 3C). This bridge also appeared at telophase (when the maternal and paternal chromatin is indistinguishable, Fig. 3E). Intriguingly, previous studies detected the same metaphase, anaphase, and telophase defects in early embryos fertilized by sperm that lack sperm-specific paternal telomere-binding proteins (*39, 41*), suggesting that maternal HipHop^yak^ interferes with paternal telomere establishment in the zygote.

We hypothesized that paternal telomere establishment, just like telomere maintenance, requires coevolution of the HipHop-HOAP interaction surface. To test this hypothesis, we again leveraged the chimeric *hiphop* genotypes (Fig. 2D and 4A). Mothers heterozygous for a *hiphop* chimera encoding the six *D. yakuba* amino acids in an otherwise *D. melanogaster* protein severely reduced progeny number (Fig. 4A). Likewise, mothers heterozygous for a *hiphop* chimera encoding the six *D. melanogaster* residues in an otherwise *D. yakuba* HipHop rescued progeny number (Fig. 4A). These data suggest that the evolution of at most six HipHop amino acids along its HOAP interaction surface is necessary and sufficient to support not only telomere maintenance through development (Fig. 2D, 2E, and S2B) but also paternal telomere establishment. Also like telomere maintenance, introducing the *D. yakuba* version of HOAP (HOAP^yak^) into the heterozygous *+/hiphop*^*yak*^ female background completely rescued their embryos’ hatch rate (Fig. S3B) and their fertility (Fig. S3C). The functional consequences of swapping the chimeric HipHop and the conspecific HOAP are consistent with coevolution of the interaction surface between these two essential telomere-binding proteins to preserve the establishment of telomere protection in the embryo.

**Figure 4.**
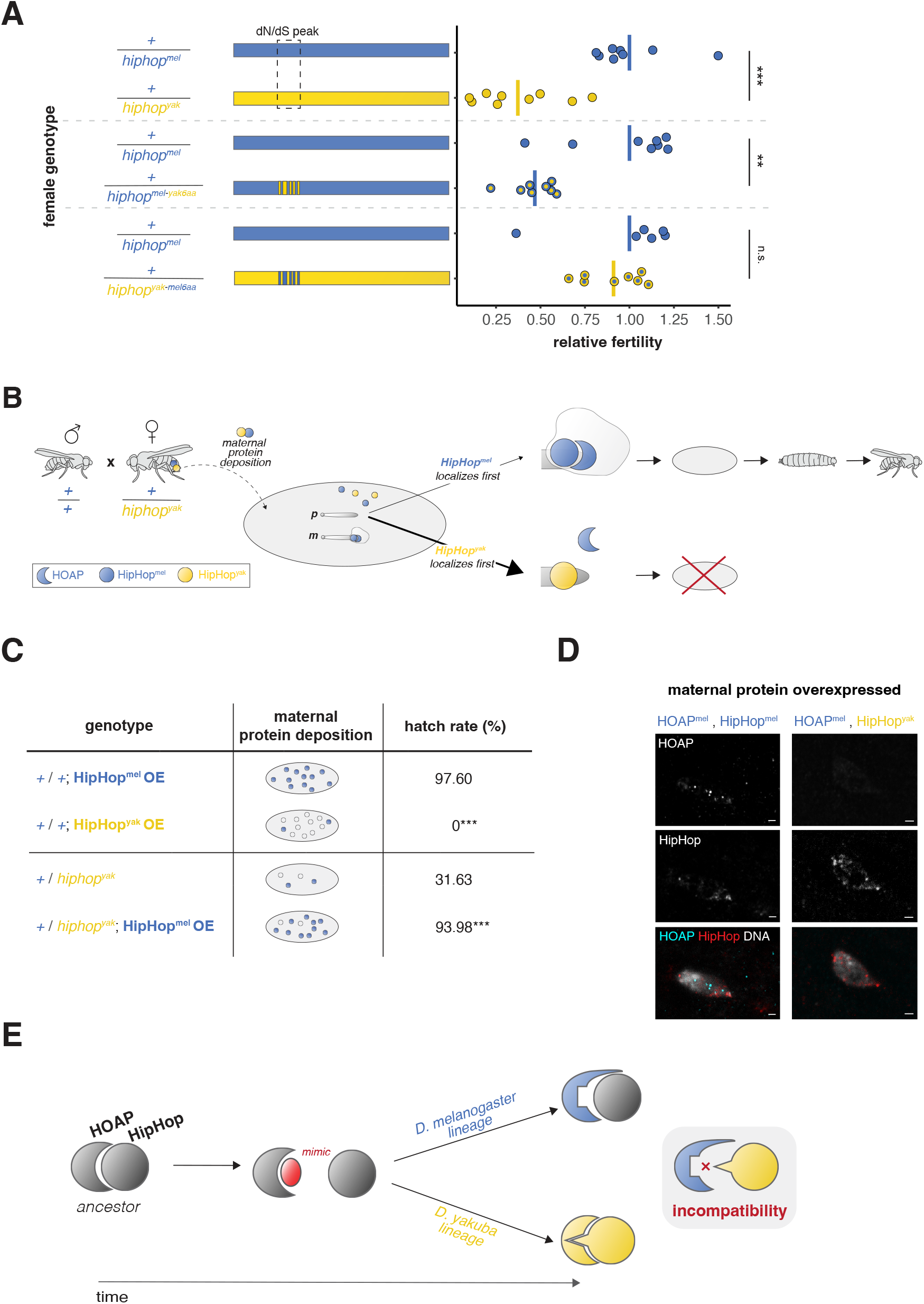
Adaptive evolution of the HipHop-HOAP interaction surface preserves telomere establishment in the early embryo. (**A**) Relative fertility of females heterozygous for *D. melanogaster*-*D. yakuba* chimeric *hiphop* transgenes crossed to wild-type males. All data points normalized by the mean fertility of *hiphop^mel^* control strain into which we introgressed each transgene. The dashed box indicates the *hiphop* coding region with elevated dN/dS. MWU test, \*\**p*-value < 0.01, *** < 0.001. (**B**) Model of dominant, maternal effect lethality induced by a single *hiphop^yak^* allele. In a small fraction of embryos from +/*hiphop^yak^* mothers, maternal HipHopmel localizes to the paternal telomere before HipHopyak and recruits endogenous HOAP. Together, the two proteins establish the paternal telomere, ensuring end-protection throughout development (upper path). In most embryos from the same mother, HipHopyak localizes to the paternal telomeres first and interferes with HOAP recruitment. The absence of HOAP inhibits the assembly of the end-protection complex, causing telomere fusions and early embryonic lethality (lower path). (**C**) Estimates of embryo hatch rate from *D. melanogaster* females overexpressing HipHopmel or HipHopyak (above) and from +/*hiphop^yak^* females with or without HipHopmel overexpression (below). n > 1100. Fisher’s Exact Test, \*\*\**p*-value < 0.001 for both comparisons. (**D**) Telophase zygotic nuclei in embryos from *D. melanogaster* females overexpressing HOAP mel and HipHopmel (left) or HOAP mel and HipHopyak (right) stained for HOAP (anti-FLAG) and HipHop (anti-EGFP). Scale bars, 1 μm. (**E**) Model of protein-protein coevolution in the end-protection complex triggered by telomeric retrotransposon evolution. A retrotransposon mimic of HipHop triggers HOAP evolution to avoid recognition of its interaction surface along both the *D. melanogaster* and *D. yakuba* lineages (Fig. S5B). Along the *D. melanogaster* lineage only, HOAP evolves adaptively at a second site to preserve interaction with HipHop, for which there is no evidence of *D*. *melanogaster*-specific, adaptive evolution (Fig. S5A). In contrast, along the *D. yakuba* lineage, HOAP evolution requires adaptive coevolution with HipHop. See Fig. S5 and S6 for additional details. “*+*” = *TM6* balancer chromosome.

We propose a model under which the specialized chromatin state of zygotic paternal chromosomes (*45-47*) combined with an initially silent zygotic genome (*44*), renders the early embryo uniquely sensitive to the presence of maternal HipHop^yak^. During normal sperm development, at least two sperm-specialized telomere-binding proteins replace the canonical end-protection proteins (*39-41*). These two proteins epigenetically mark paternal telomeres for recognition and replacement by the maternally provisioned, canonical end-protection proteins in the embryo (*39*). One such sperm-specific telomere-binding protein, K81, is encoded by a young duplicate gene of *hiphop* (*39, 48*). Maternal HipHop replaces paternal K81 at fertilization (*39*) and recruits other maternally provisioned members of the canonical end-protection complex to paternal telomeres. Mothers heterozygous for *hiphop*^*yak*^ provision their embryos with both Hiphop^yak^ and the endogenous HipHop^mel^. These embryos’ high rate of lethality (Fig. 1E), combined with the aberrantly elevated signal of Hiphop^yak^ at *D. melanogaster* telomeres (Fig. 2B), predicts that maternally provisioned HipHop^yak^ outcompetes HipHop^mel^ in the replacement of K81 at paternal telomeres (Fig. 4B). Under this model, the binding of HipHop^yak^ to paternal telomeres prevents HOAP recruitment and, consequently, the assembly of the essential end-protection complex. A small fraction of embryos escape and survive to adulthood. In these escaper embryos, we predict that HipHop^mel^ localizes to the telomere before HipHop^yak^. HipHop^mel^ recruits endogenous HOAP, preserving paternal telomere stability and early embryogenesis. The presence of at least one zygotic copy of the endogenous *D. melanogaster hiphop* enables the epigenetic propagation of the end-protection complex through development (Fig. 4B).

To test this “competition model” of telomere establishment, we manipulated the maternal dose of the conspecific and heterospecific versions of HipHop deposited into the *D. melanogaster* embryo. The model predicts that elevating the maternal dose of HipHop^yak^ would elevate the fraction of embryos that die in a *D. melanogaster* background. Consistent with this prediction, overexpressing HipHop^yak^ in *D. melanogaster* mothers resulted in completely penetrant embryonic lethality (Fig. 4C). Likewise, we expected to mitigate embryonic lethality caused by the single *hiphop*^*yak*^ allele upon elevating the dose of HipHop^mel^. As predicted, we observed a 3-fold higher hatch rate in the embryos from +/*hiphop*^*yak*^ mothers that overexpress HipHop^mel^ (Fig. 4C). These data are consistent with HipHop^yak^ outcompeting HipHop^mel^ for localization to the paternal telomere and so preventing HOAP recruitment and telomere establishment. Further consistent with the competition model, we observed HOAP^mel^ and HipHop^mel^ localization to embryonic telophase chromosomes in the HipHop^mel^-deposited embryos, but we failed to detect HOAP on telophase chromosomes in HipHop^yak^-deposited embryos (Fig. 4D).

Our data are consistent with a single mechanism underlying this dominant maternal-effect lethality and the recessive larval lethality: HipHop^yak^ disrupts endogenous HOAP localization to the telomere and, consequently, the assembly of the end-protection complex. In the zygotes of +/*hiphop*^*yak*^ heterozygous mothers, *hiphop*^*yak*^ acts dominantly because maternal HipHop^yak^ outcompetes maternal HipHop^mel^ for telomere localization (Fig. 4B). HipHop^yak^ prevents HOAP localization, which compromises end protection and causes lethality in most embryos. In rare cases, maternal HipHop^mel^ localizes first and establishes telomere end-protection. If the “escapers” are heterozygous for *hiphop*^*yak*^, the single zygotic copy of *D. melanogaster hiphop* is sufficient to maintain the integrity of the telomere complex in the larva, pupa, and adult. That is, once maternal HipHop^mel^ establishes proper end-protection, the complex is epigenetically propagated and sustained by HipHop^mel^, even in the presence of HipHop^yak^ (Fig. S4A). Indeed, overexpression of HipHop^yak^ after telomere establishment in an otherwise wild-type *D. melanogaster* has no effect on viability (Fig. S4B). If the escaper embryos are instead homozygous for *hiphop*^*yak*^, lethal telomere fusions accumulate as the maternally provisioned HipHop^mel^ (from *+/hiphop*^*yak*^ heterozygous mothers) depletes through development, leaving only zygotic HipHop^yak^ in the late larval stage. The contrasting dominant maternal-effect lethality in the embryo and recessive lethality in the larva suggest that the highly specialized chromatin state of sperm-deposited paternal DNA renders the zygote uniquely vulnerable to the evolution of chromosomal proteins.

Our study of HipHop-HOAP coevolution demonstrates that preserving an ostensibly conserved, essential function, chromosome end-protection, requires recurrent evolutionary innovation of a protein-protein interaction surface. The short evolutionary distance between *D. melanogaster* and *D. yakuba* enabled us to deconvolve, for the first time, the often confounded effects of neutral and adaptive evolution in gene swaps between distantly related species (*22, 23*). Indeed, the short evolutionary distance and our statistical analyses (Table S1 and S2) implicate adaptive, rather than neutral, evolution. Adaptive evolution implicates an ever-changing evolutionary pressure on telomere-binding proteins to change. Previous research suggests that fast-evolving, selfish retrotransposons impose a strong evolutionary pressure on this complex. HOAP evolves adaptively to restrict telomeric retrotransposon insertions (*42*) and HipHop promotes silencing of these retrotransposons (*49*). Combined with evidence that the dose of these telomere end-protection proteins determines retrotransposon activity (*42, 49*), we speculate that a telomere-enriched selfish retrotransposon in *Drosophila* encodes a mimic (*50, 51*) of the HipHop-HOAP interaction surface that partially destabilizes the complex (Fig. 4E, S5, and S6). In response, HOAP evolves to avoid recognition by altering both common and distinct sites along the *D. melanogaster* and *D. yakuba* lineages (Fig. S5B). The restriction of HipHop adaptive evolution to the *D. yakuba* branch (Fig. S5A) suggests distinct, lineage-specific coevolutionary dynamics in response to this mimic (Fig. 4E and S6). Importantly, the observation that females heterozygous for *hiphop*^*mel*^ and *hiphop*^*yak*^ manifest this inter-species incompatibility offers a new mechanism for how conflict with selfish genetic elements can trigger F_1_ hybrid sterility, a classic reproductive barrier between species (*52, 53*).

Our study also highlights the hidden complexity of adaptive coevolution within multi-protein complexes. The discovery that a single copy of a heterologous allele is sufficient to cause lethality, i.e., dominant maternal effect lethality, rejects the arguably more intuitive prediction that disrupting an interaction surface would result in a recessive, loss-of-function phenotype (*24, 43*). Also unexpectedly, we discovered that protein interdependence itself may evolve – while HipHop depends on HOAP for telomere localization in *D. melanogaster*, the *D. yakuba* version of HipHop localized to telomeres independently of its interaction partner. These distinct mechanisms of telomere protein recruitment implicate distinct evolutionary paths to preserving an essential function. These advances highlight how leveraging *in vivo*, experimental dissection of coevolution is vital to understanding how adaptive evolution both perturbs and preserves essential, multi-protein complex integrity. As we enter this era of increasing genomic sequence data and AI-based protein structure prediction tools (*54-58*), many candidate coevolving proteins will emerge. We anticipate that future experimental manipulations of coevolving partners will reveal the multitude of ways adaptive evolution tweaks even the most conserved biological functions.

## Materials and Methods

### Molecular evolution and population genetic analyses

To test for positive selection across the end-protection complex, we conducted a phylogeny-based molecular evolution analysis using orthologs of the *melanogaster* subgroup (*D. melanogaster, D. simulans, D. sechellia, D. mauritiana, D. erecta, D. yakuba*, and *D. teissieri*, Table S3). We aligned nucleotide and protein sequences of each gene using MUSCLE (v5.1) (*59*) and generated PHYLIP files using PAL2NAL (*60*). For each gene, we fit the multiple alignment to an NSsites model using the codeml program in the PAML software package (v4) (*61*). We compared the likelihood of models M7 (neutral: the dN/dS values fit a beta distribution between 0 and 1) and M8 (non-neutral: M7 parameters plus dN/dS > 1) using the F3×4 model of codon frequency using a x^2^ test.

To test for positive selection at the *hiphop* locus using *D. melanogaster* and *D. yakuba* only, we performed a McDonald-Kreitman (*62*) test followed by Fisher’s Exact Test. This test detects adaptive evolution by comparing synonymous and nonsynonymous polymorphisms within species to synonymous and nonsynonymous fixations between species. To generate these counts, we extracted 194 *D. melanogaster* alleles of *hiphop*, including one allele from the reference genome (dmel r6.53) and 193 alleles from the genomes of Zambian *D. melanogaster* strains (DPGP3, Table S3) (*63*). We extracted the *D. yakuba* allele from the reference genome assembly (Prin_Dyak_Tai18E2_2.1). We aligned the nucleotide and protein sequences using MUSCLE (*59*) and generated a final alignment of in-frame codons using PAL2NAL (*60*). To identify HipHop domains enriched for signals of positive selection, we estimated the dN/dS of 50-base pair (bp) windows, sliding 10-bp along an alignment of the *D. melanogaster* and *D. yakuba* coding sequences (dmel r6.54 and Prin_Dyak_Tai18E2_2.1, respectively, DnaSP v6 (*64*)).

To assess lineage-specific molecular evolution of *hiphop*, we first reconstructed the ancestral *hiphop* allele of *D. melanogaster* and *D. yakuba* using *D. erecta* as an outgroup to polarize changes (NSsites model M0 from the codeml program in PAML (v4) (*61*)). We then estimated pairwise dN/dS using the reconstructed ancestor and either the *D. melanogaster* or *D. yakuba hiphop* allele (window size of 100 bp, 20 bp slide).

### Fly stock construction

#### Whole gene and chimeric gene swaps

We used CRISPR/Cas9 to generate *D. melanogaster* flies that encode an EGFP-tagged, codon-optimized version of either the *D. melanogaster* coding sequence or the *D. yakuba* coding sequence of *hiphop*, both swapped into the endogenous *hiphop* locus. We first generated a U6.2 promoter-driven guide RNA construct by cloning sgRNAs flanking the coding sequence of *hiphop* (upstream: 5’-GGTGCATGATCTATTTCAGA-3’, downstream: 5’-GTACTTGATGGGAACCACAGG-3’) into pBFv-U6.2 and pBFv-U6.2B backbones (*65*). We shuttled the downstream sgRNA into pBFv-U6.2 to create a dual sgRNA vector (University of Utah Mutagenesis Core). In parallel, we constructed homology-directed repair (HDR) plasmids (*66*) encoding one kilobase homology arms 5’ and 3’ of their respective guide RNAs. Between the homology arms, we synthesized a codon-optimized (for *D. melanogaster*) *hiphop* coding sequence from either *D. melanogaster* or *D. yakuba* (GenScript, Piscataway, NJ). We N-terminally tagged each sequence with EGFP separated by a linker sequence (GGTGGTTCATCA). We injected the dual sgRNA vector and a single HDR plasmid into Cas9-expressing lines, vas-Cas9 (X) RFP+ GFP+ (BDSC# 51323) and yw; nos-Cas9 (II-attP40) for *D. melanogaster* and *D. yakuba hiphop* transgenic lines, respectively (BestGene, Chino Hills, CA).

To screen for transformants, we crossed the G_0_ adults, injected as embryos, to a *w; Pin/CyO; Dr/TM6B* stock. We screened F_1_ females to identify positive transformants using forward primer 5’-GTGGCGGGACAATTGGCTTCATG-3’ and reverse primer 5’-TCCTCGATGTTGTGGCGGATC-3’. We then generated long-term stocks of either *hiphop*^*mel*^ or *hiphop*^*yak*^ allele. To confirm that positive transformants encoded a complete transgene in the correct genomic location, we amplified the gene region from homozygous *hiphop*^*mel*^ and heterozygous *hiphop*^*yak*^ flies (homozygous *hiphop*^*yak*^ flies are lethal) using primers that anneal outside of the homology arms (5’-AGACACTCACCAACACCAGCA-3’ and 5’-TAAGAGGGCACGATTCCACA-3’) and sequenced across the entire region (Table S4). For heterozygous *hiphop*^*yak*^ flies, we distinguished *hiphop*^*yak*^ from the endogenous *D. melanogaster hiphop* based on different amplicon sizes. We then sequenced the gel-extracted product of *hiphop*^*yak*^ (Table S4). We also designed primers that amplified the endogenous *hiphop* locus (5’-GTGGCGGGACAATTGGCTTCATG-3’, 5’-CTGGTAGTGGGCACATTGTCTGAATCTTG-3’) to confirm that our *hiphop*^*mel*^ was a true replacement.

To generate the *hiphop*^*yak*^,*HOAP*^*yak*^ recombinant chromosome, we crossed *hiphop*^*yak*^ to a previously constructed CRISPR/Cas9 *D. melanogaster* transgenic fly encoding the *D. yakuba* version of *HOAP* (*42*). We crossed these F_1_ females to *w/Y; +/+; TM3/TM6B* males and screened single F_2_ males to identify the desired *hiphop*^*yak*^,*HOAP*^*yak*^ recombinant chromosome using primers 5’-GACCAGCCCTTTATTGACATTTACT-3’ and 5’-AAGTTGTGAACCTCTTCCAG-3’ for *hiphop*^*yak*^ and primers 5’-CAAATGGACCCACCAATTCCGAGAG-3’ and 5’-TCACCGTCATGGTCTTTGTAGTCCAT-3’ for *HOAP*^*yak*^. We then backcrossed positive F_2_ males to *w; +/+; TM3/TM6B* females and self-crossed the balanced F_3_ progeny to generate the *hiphop*^*yak*^,*HOAP*^*yak*^ stock.

To generate the *D. melanogaster-D. yakuba* chimeric *hiphop* transgenic lines in *D. melanogaster*, we used CRISPR/Cas9 to replace the endogenous *hiphop* locus with either a *hiphop*^*mel-yak6aa*^ allele or a *hiphop*^*yak-mel6aa*^ allele (WellGenetics Inc., New Taipei, Taiwan). We first cloned sgRNAs flanking the coding sequence of *hiphop* (upstream: 5’-GGTGCATGATCTATTTCAGA-3’, downstream: 5’-TACTTGATGGGAACCACAGG-3’) downstream of a U6 promoter on the pDCC6 plasmid (*67*). In parallel, we constructed HDR plasmids with the *egfp-chimeric hiphop-PBacDsRed* cassette flanked with one kilobase homology arms 5’ and 3’ of their respective guide RNAs into pUC57-Kan (GenScript, Piscataway, NJ). The cassette consists of the *3xP3-DsRed* visible marker (*66*) flanked by PBac transposon ends (https://flycrispr.org/scarless-gene-editing/) and encodes an N-terminally EGFP-tagged chimeric coding sequences of either *hiphop*^*mel*^ with *D. yakuba* amino acids 36H, 39W, 40R, 47V, 49Q, and 58H (*hiphop*^*mel-yak6aa*^) or *hiphop*^*yak*^ with *D. melanogaster* amino acids 36R, 39C, 40L, 47A, 49N, and 58D (*hiphop*^*yak-mel6aa*^) with a linker GGTGGATCCTCA (also codon-optimized for *D. melanogaster*). We injected the pDCC6 plasmid (*67*) containing Cas9 and sgRNAs and the HDR plasmid into the *w*^*1118*^ stock and selected positive F_1_ flies carrying the visible marker 3xP3-DsRed. We sequence-validated positive transformants with primer pairs that annealed outside the homology arm (upstream: 5’-CAGACAACGAGTCAGACACACA-3’ and 5’-GAACTTCAGGGTCAGCTTGC-3’, downstream: 5’-TTTGACTCACGCGGTCGTTA-3’ and 5’-AGAGAGGCGGCTTTTGAACT-3’). We then excised the visible marker 3xP3-DsRed by PiggyBac transposition. We sequence-validated these excision lines using primers 5’-CCCGACAACCACTACCTGAG-3’ and 5’-AGTTGTGAACCTCTTCCAGTG-3’. To enable rigorous fertility comparisons across *hiphop*^*mel*^, *hiphop*^*mel-yak6aa*^, and *hiphop*^*yak-mel6aa*^, each of which was engineered independently, we introgressed each transgenic chromosome into the *hiphop*^*mel*^ stock using balancer chromosomes.

#### Overexpression genotypes

We used the *<!C31* integrase-mediated transgenesis system (*68*) to construct chromosomes with a *UASp-egfp-hiphop*^*mel*^ or *UASp-egfp-hiphop*^*yak*^ transgene into the same attP landing site (attP40). We first PCR-amplified the *egfp-hiphop* coding sequences (either the *D. melanogaster* or *D. yakuba* allele) using Phusion High-Fidelity DNA Polymerase (NEB, Ipswich, MA) from the HDR plasmids (see above). We cloned the PCR products into *NotI/BamHI* sites of the pUASp-attB vector (*Drosophila* Genomics Resource Center, Bloomington, IN) and validated with Sanger sequencing. We then introduced the constructs into *D. melanogaster y*^*1*^ *w*^*67c23*^; *P{CaryP}attP40* flies, which have an attP landing site at cytological position 25C6 on chromosome *2L* (The BestGene, Inc., Chino Hills, CA).

To overexpress the transgenic *hiphop* alleles ubiquitously, we crossed each UAS-transgene stock to *yw; +/+; Act5C-GAL4/TM6B* (BDSC #3954). To overexpress *hiphop* alleles in the female germline, we crossed each UAS-transgene stock to nos-*GAL4-VP16* (BDSC #64277). To assess fertility rescue of +/*hiphop*^*yak*^ by overexpressing *hiphop*^*mel*^, we crossed *w; UASp-hiphop*^*mel*^*/Gla; nos-GAL4-VP16* to *w; +/+; hiphop*^*yak*^*/TM6B*. We assayed fertility of genotypes *w; Gla/+; nos-GAL4-VP16 /hiphop*^*yak*^ (control) and *w; UASp-hiphop*^*mel*^*/+; nos-GAL4-VP16 /hiphop*^*yak*^ (experiment).

To overexpress the *D. melanogaster HOAP*, we used the *<!C31* integrase-mediated transgenesis system (*68*) to construct a chromosome with a *UASp-HOAP*^*mel*^*-FLAG* transgene. We first synthesized a *3xFLAG*-tagged *D. melanogaster HOAP* coding sequence and cloned it into the pUASp-attB vector (Twist Bioscience, South San Francisco, CA). We then introduced the constructs into *D. melanogaster y*^*1*^ *w*^*67c23*^; *P{CaryP}attP40*, which have an attP transgene landing site at cytological position 25C6 on chromosome *2L* (BestGene, Chino Hills, CA). We used this transgene to increase *HOAP*^*mel*^ maternal deposition into embryos along with HipHop^mel^ or HipHop^yak^ to detect HOAP and HipHop, the endogenous versions of which are below the detection limit. We assayed progeny of the flies with genotypes *w; UASp-hiphop*^*mel*^*/UASp-HOAP*^*mel*^; *nos-GAL4-VP16/+* or *UASp-hiphop*^*yak*^*/UASp-HOAP*^*mel*^; *nos-GAL4-VP16/+*.

### Fitness assays

#### Viability

To assay adult viability, we collected virgin females and males from *hiphop*^*mel*^*/TM6B, hiphop*^*yak*^*/TM6B, hiphop*^*yak-mel6aa*^*/TM6B, hiphop*^*mel-yak6aa*^*/TM6B*, or *hiphop*^*yak*^,*HOAP*^*yak*^*/TM6B*. For each replicate vial, we crossed three virgin females to three males. For *hiphop*^*yak*^ and *hiphop*^*mel-*^ _*yak6aa*_, which have severely reduced female fertility in the heterozygous state, we instead crossed six virgin females to six males. We flipped the parents onto new food every three days for 12 days and scored adult progeny as homozygous or heterozygous. We normalized the percent homozygous adults by the mean percent of *hiphop*^*mel*^ homozygous adults. We then performed Mann-Whitney U test to compare mean differences.

To assay pupal viability, we collected virgin females and males carrying the *hiphop*^*mel*^/*TM6B* or *hiphop*^*yak*^/*TM6B* heterozygous genotypes. For each replicate vial, we crossed three virgin females to three males for *hiphop*^*mel*^ or six virgin females to six males for *hiphop*^*yak*^. We flipped the parents onto new food every three days for six days and assessed the number of pupae homozygous and heterozygous for *hiphop*. We normalized the percent *hiphop*^*yak*^ homozygotes by the percent *hiphop*^*mel*^ homozygotes. We then performed Mann-Whitney U test to compare mean differences.

To assay adult viability of flies ubiquitously overexpressing either *hiphop*^mel^ or *hiphop*^yak^, we crossed either three *UASp-hiphop*^*mel*^ or three *UASp-hiphop*^*yak*^ virgin females to three *yw; +/+; Act5C-GAL4/TM6B* males in each replicate vial. We flipped the parents onto new food every three days for 12 days and scored all progeny that emerged carrying *Act5C-GAL4* or *TM6B*. We estimated the percent *UASp-hiphop*^*mel*^*/+; Act5C-GAL4/+* progeny and *UASp-hiphop*^*yak*^*/+; Act5C-GAL4/+* progeny and normalized to the former. We then performed Mann-Whitney U test to compare mean differences.

#### Fertility, fecundity, and hatch rate assays

To assay female fertility of *hiphop*^*mel*^, *hiphop*^*yak*^, *hiphop*^*yak-mel6aa*^, *hiphop*^*mel-yak6aa*^, *hiphop*^*yak*^,*HOAP*^*yak*^, and the *hiphop* mutant *P{hsneo}hiphop*^*1*^, we first collected 1-5-day-old, heterozygous virgin females over *TM6B*. For each replicate vial, we crossed three virgin females to three *w*^*1118*^ males. We flipped the parents onto new food every three days for 12 days and counted all progeny that emerged. To assay male fertility of *hiphop*^*mel*^ and *hiphop*^*yak*^ heterozygotes over *TM6B*, we first collected 1-5-day-old *w*^*1118*^ females. For each replicate vial, we crossed three *w*^*1118*^ virgin females to three *hiphop*^*mel*^ or *hiphop*^*yak*^ heterozygous males. We flipped the parents onto new food every three days for 12 days and counted all progeny that emerged. We then performed Mann-Whitney U test to compare mean differences.

To assay the number of mature eggs produced by *hiphop*^*mel*^ or *hiphop*^*yak*^ heterozygous females over *TM6B*, we counted eggs with elongated dorsal appendages from 3–5-day-old females. We then performed Mann-Whitney U test to compare mean differences.

To conduct hatch rate assays, we let intercrossed *hiphop*^*mel*^, *hiphop*^*yak*^, or *hiphop*^*yak*^,*HOAP*^*yak*^ heterozygous females over *TM6B* and females overexpressing *hiphop* alleles in the female germline lay for two hours after 1-hour pre-lay. We counted the total number of embryos and the number of larvae that hatched daily over a 72-hour window. We then performed Fisher’s Exact Test to assess deviations from the null expectation.

We conducted all assays on molasses food or Nutri-Fly^®^ Grape Agar media (Genesee Scientific, Morrisville, NC) at 25°C.

### Immunofluorescence

#### chromosomes from larval brains

To assess telomere fusion rates and telomere protein localization, we dissected the brains from third instar larvae, the developmental stage immediately before lethality of the homozygous *hiphop*^*yak*^ genotype. We dissected in a saline solution (0.7% NaCl) and then incubated in 1.5 × 10^−5^ M colchicine for 1 hour to enrich for metaphase chromosomes. To promote chromosome spreading, we treated the brains with a hypotonic solution (0.5% sodium citrate) for 10 minutes. We then placed the samples in a fixative solution (1.8% paraformaldehyde in 45% acetic acid) for 20 minutes. Following fixation, we squashed the brains in the fixative solution on poly-L-lysine coated glass slides (Sigma-Aldrich, St. Louis, MO) using coverslips treated with Sigmacote (Sigma-Aldrich, St. Louis, MO). After squashing, we flash-froze the samples in liquid nitrogen and flicked off the coverslips with a razor blade. We then transferred the slides into 100% ethanol for 10 minutes at -20°C. We next washed the slides twice in 1x PBS for 10 minutes each and twice in 1x PBS, 0.1% Tween 20 (1x PBST) (Sigma-Aldrich, St. Louis, MO) for 10 minutes each. After washing, we blocked the slides in 1x PBST, 3% BSA for 1 hour. We then incubated the slides with mouse anti-FLAG M2 (Sigma-Aldrich, St. Louis, MO, 1:2500) and chicken anti-GFP (Aves Labs, Davis, CA,1:500), diluted in blocking solution, overnight at 4°C. We then washed the slides three times in 1x PBST, 1% BSA for 10 minutes each and then incubated for 2 hours at room temperature in anti-mouse Alexa Fluor 488 and anti-chicken Alexa Flour 568 (Thermo Fisher Scientific, Waltham, MA, 1:300), diluted in block solution. We then washed the slides in 1x PBST with 3% BSA, 1x PBST with 1% BSA, 1x PBST, and 1x PBS for 10 minutes each. Finally, we mounted the samples with SlowFade Gold Antifade Mountant with DAPI (Thermo Fisher Scientific, Waltham, MA).

#### Embryos

After a 1-hour pre-lay, we collected embryos for 40 minutes from *hiphop*^*mel*^*/TM6B* and *hiphop*^*yak*^*/TM6B* females and for 1 hour from *w; UASp-HOAP*^*mel*^*-FLAG/UASp-hiphop*^*mel*^*-EGFP; nos-GAL4-VP16/+*, and *UASp-HOAP*^*mel*^*-FLAG/UASp-hiphop*^*yak*^*-EGFP; nos-GAL4-VP16/+* females (*69*). We then fixed each genotype separately using methanol and heptane (*70*) and immunostained following Loppin *et al*. (*71*) (rabbit anti-AcH4, MilliporeSigma, Burlington, MA, 1:1000, mouse anti-FLAG M2, Sigma-Aldrich, St. Louis, MO, 1:500, and chicken anti-GFP, Aves Labs, Davis, CA, 1:500). We used the following secondary antibodies: goat anti-rabbit Alexa Flour 568 for the 40-minute collection and goat anti-mouse Alexa Fluor 488, goat anti-chicken Alexa Flour 568, and goat anti-rabbit Alexa Flour 633, for the 1 hour collection (all from Thermo Fisher Scientific, Waltham, MA at 1:500).

### Determining embryonic nuclear cycle number

Zygotic genome activation (ZGA) occurs at embryonic nuclear cycle 14, about 2.5 hours post-fertilization. To determine the percent of embryos that develop beyond ZGA, we self-crossed *hiphop*^*mel*^*/TM6B* or *hiphop*^*yak*^*/TM6B*. We collected embryos for one hour and then aged the collection for 2.5 hours (after 1-hour pre-lay). We then used methanol and heptane to fix each genotype separately (*70*) and mounted the samples with SlowFade Gold Antifade Mountant with DAPI (Thermo Fisher Scientific, Waltham, MA). We calculated the percent embryos with nuclei at the periphery (rather than in the interior), a marker of ZGA (*72*). We then performed Fisher’s Exact Test between the two genotypes.

To assess developmental progression in even earlier embryos, we allowed the same heterozygous females (crossed to heterozygous males) to lay for 1 hour after a 1-hour pre-lay. We then methanol-fixed each genotype separately (*70*) and immunostained with AcH4 (rabbit anti-AcH4, MilliporeSigma, Burlington, MA, 1:1000, following Loppin et al. (*71*)) and mounted the embryos as described above. For this particular experiment, AcH4 enabled us to delineate pronuclear migration (when only the paternal chromosomes are positive) from nuclear cycle 2 (when both maternal and paternal chromosomes are positive). We then calculated percent embryos at each nuclear cycle 1-7 and performed Fisher’s Exact Test between the two genotypes.

### Fluorescence *in situ* hybridization

We dissected third instar larval salivary glands into 1x PBS, 0.1% Tween 20 (1x PBST) and fixed in 2% paraformaldehyde in 45% acetic acid for 1 minute. Following fixation, we placed the salivary glands in 45% acetic acid between poly-L-lysine coated glass slides (Sigma-Aldrich, St. Louis, MO) and coverslips treated with Sigmacote (Sigma-Aldrich, St. Louis, MO). We then gently moved the coverslip back and forth to facilitate cell lysis and chromosome spreading and squashed with a rubber hammer. We next flash-froze the samples in liquid nitrogen and flicked off the coverslips with a razor blade. We then transferred the slides into 100% ethanol for 10 minutes at -20°C and followed the *in situ* hybridization protocol described in Saint-Leandre *et al*. (*42*) using a probe cognate to the HeT-A retrotransposon (see Table S4).

### Analysis of cytological data

We calculated telomere fusion rates by dividing counts of mitotic cells with three or more telomere associations by the total number of mitotic cells (>100 mitotic cells per individual larva, ≥ three larvae per genotype for the whole gene swaps and ;:58 mitotic cells from two individuals for the chimeric gene swaps). We combined data from all individuals per genotype and performed Fisher’s Exact Test. We quantified HOAP signal at ≥165 chromosome ends per genotype and HipHop signal at ≥87 chromosome ends per genotype from at least three mitotic cells from three individual larvae. We then performed Mann-Whitney U to compare mean differences. To assess maternal-paternal chromosome asymmetry at metaphase, we calculated a circularity index 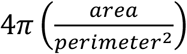(FIJI software, (*73*)) of the AcH4-positive and negative chromosomes (paternal and maternal, respectively). We then performed Mann-Whitney U test to compare mean differences.

We imaged all slides on a Leica TCS SP8 Four Channel Spectral Confocal System. For each experiment, we used the same imaging parameters for all genotypes.

## Supporting information

LinETAL_Supplement

## Acknowledgments

The authors thank members of the Levine Lab, Lampson Lab, H. Malik, N. Elde, A. Das, Y. Rong, and B. Loppin for vital discussions about the project. The authors also thank M. Lampson, C. Brand, and D. Dudka for feedback on earlier versions of the manuscript and S. Mohammed for technical assistance. The University of Utah Mutation Generation and Detection Core provided assistance in developing the guide RNA expression constructs.

## Funding

Taiwanese Government Scholarship for Study Abroad to S.L. and National Institutes of Health grant R35GM124684 to M.T.L.

## Author contributions

M.T.L. conceived the study. S.L. and M.T.L. designed and led the project. S.L and H.F. conducted the experiments. S.L. analyzed the data. S.L. and M.T.L drafted the manuscript. All authors contributed revisions and approved the final version of the manuscript.

## Competing interests

The authors declare no competing interests.

## Data and materials availability

All transgenic stocks are available upon request and all raw data will be deposited into Dryad Digital Repository.

## Notes

### Competing Interest Statement

The authors have declared no competing interest.

## Literature Cited

1. S. Lindquist, E. A. Craig, The heat-shock proteins. Annu Rev Genet 22, 631–677 (1988).

2. N. C. Elde, S. J. Child, A. P. Geballe, H. S. Malik, Protein kinase R reveals an evolutionary model for defeating viral mimicry. Nature 457, 485–489 (2009).

3. H. S. Malik, S. Henikoff, Adaptive evolution of Cid, a centromere-specific histone in Drosophila. Genetics 157, 1293–1298 (2001).

4. M. G. Schueler, W. Swanson, P. J. Thomas, N. C. S. Program, E. D. Green, Adaptive evolution of foundation kinetochore proteins in primates. Mol Biol Evol 27, 1585–1597 (2010).

5. Y. C. Lee, C. Leek, M. T. Levine, Recurrent innovation at genes required for telomere integrity in Drosophila. Mol Biol Evol 34, 467–482 (2017).

6. C. N. Passow et al., Contrasting patterns of rapid molecular evolution within the p53 network across mammal and sauropsid lineages. Genome Biol Evol 11, 629–643 (2019).

7. D. C. Presgraves, W. Stephan, Pervasive adaptive evolution among interactors of the Drosophila hybrid inviability gene, Nup96. Mol Biol Evol 24, 306–314 (2007).

8. P. A. Rowley, K. Patterson, S. B. Sandmeyer, S. L. Sawyer, Control of yeast retrotransposons mediated through nucleoporin evolution. Plos Genetics 14, (2018).

9. S. C. Lovell, D. L. Robertson, An integrated view of molecular coevolution in proteinprotein interactions. Mol Biol Evol 27, 2567–2575 (2010).

10. A. K. Ramani, E. M. Marcotte, Exploiting the co-evolution of interacting proteins to discover interaction specificity. J Mol Biol 327, 273–284 (2003).

11. C. S. Goh, A. A. Bogan, M. Joachimiak, D. Walther, F. E. Cohen, Co-evolution of proteins with their interaction partners. Journal of Molecular Biology 299, 283–293 (2000).

12. R. Jothi, P. F. Cherukuri, A. Tasneem, T. M. Przytycka, Co-evolutionary analysis of domains in interacting proteins reveals insights into domain-domain interactions mediating protein-protein interactions. Journal of Molecular Biology 362, 861–875 (2006).

13. S. Salmanian, H. Pezeshk, M. Sadeghi, Inter-protein residue covariation information unravels physically interacting protein dimers. BMC Bioinformatics 21, 584 (2020).

14. L. Rosin, B. G. Mellone, Co-evolving CENP-A and CAL1 domains mediate centromeric CENP-A deposition across Drosophila species. Dev Cell 37, 136–147 (2016).

15. S. Yang, J. Wang, R. T. Ng, Inferring RNA sequence preferences for poorly studied RNA-binding proteins based on co-evolution. BMC Bioinformatics 19, 96 (2018).

16. R. M. Strange, L. P. Russelburg, K. J. Delaney, Co-evolution of SNF spliceosomal proteins with their RNA targets in trans-splicing nematodes. Genetica 144, 487–496 (2016).

17. S. Yang et al., Correlated evolution of transcription factors and their binding sites. Bioinformatics 27, 2972–2978 (2011).

18. M. A. Beilstein et al., Evolution of the telomere-associated protein POT1a in Arabidopsis thaliana is characterized by positive selection to reinforce protein-protein interaction. Molecular Biology and Evolution 32, 1329–1341 (2015).

19. D. de Juan, F. Pazos, A. Valencia, Emerging methods in protein co-evolution. Nat Rev Genet 14, 249–261 (2013).

20. F. Morcos, J. N. Onuchic, The role of coevolutionary signatures in protein interaction dynamics, complex inference, molecular recognition, and mutational landscapes. Curr Opin Struct Biol 56, 179–186 (2019).

21. J. Durham, J. Zhang, I. R. Humphreys, J. Pei, Q. Cong, Recent advances in predicting and modeling protein-protein interactions. Trends Biochem Sci 48, 527–538 (2023).

22. H. Y. Lai, Y. H. Yu, Y. T. Jhou, C. W. Liao, J. Y. Leu, Multiple intermolecular interactions facilitate rapid evolution of essential genes. Nat Ecol Evol 7, 745–755 (2023).

23. H. Takeuchi, S. Nagahara, T. Higashiyama, F. Berger, The chaperone NASP contributes to de novo deposition of the centromeric histone variant CENH3 in Arabidopsis early embryogenesis. Plant Cell Physiol 65, 1135–1148 (2024).

24. I. Sandler, M. Abu-Qarn, A. Aharoni, Protein co-evolution: How do we combine bioinformatics and experimental approaches? Mol Biosyst 9, 175–181 (2013).

25. B. M. Moran et al., A lethal mitonuclear incompatibility in complex I of natural hybrids. Nature 626, 119–127 (2024).

26. W. El Yakoubi, T. Akera, Condensin dysfunction is a reproductive isolating barrier in mice. Nature 623, 347–355 (2023).

27. K. B. S. Swamy et al., Proteotoxicity caused by perturbed protein complexes underlies hybrid incompatibility in yeast. Nat Commun 13, 4394 (2022).

28. A. Lukacs et al., The Integrity of the HMR complex is necessary for centromeric binding and reproductive isolation in Drosophila. PLoS Genet 17, e1009744 (2021).

29. M. Kitaoka, O. K. Smith, A. F. Straight, R. Heald, Molecular conflicts disrupting centromere maintenance contribute to Xenopus hybrid inviability. Curr Biol 32, 3939–3951 e3936 (2022).

30. G. D. Raffa, L. Ciapponi, G. Cenci, M. Gatti, Terminin: A protein complex that mediates epigenetic maintenance of Drosophila telomeres. Nucleus 2, 383–391 (2011).

31. L. Cheng, M. Cui, Y. S. Rong, MTV sings jubilation for telomere biology in Drosophila. Fly (Austin) 12, 41–45 (2018).

32. G. Cenci, G. Siriaco, G. D. Raffa, R. Kellum, M. Gatti, The Drosophila HOAP protein is required for telomere capping. Nat Cell Biol 5, 82–84 (2003).

33. G. Gao et al., HipHop interacts with HOAP and HP1 to protect Drosophila telomeres in a sequence-independent manner. EMBO J 29, 819–829 (2010).

34. Y. Zhang et al., MTV, an ssDNA protecting complex essential for transposon-based telomere maintenance in Drosophila. PLoS Genet 12, e1006435 (2016).

35. G. D. Raffa et al., The Drosophila modigliani (moi) gene encodes a HOAP-interacting protein required for telomere protection. Proc Natl Acad Sci U S A 106, 2271–2276 (2009).

36. G. D. Raffa et al., Verrocchio, a Drosophila OB fold-containing protein, is a component of the terminin telomere-capping complex. Genes Dev 24, 1596–1601 (2010).

37. L. Fanti, G. Giovinazzo, M. Berloco, S. Pimpinelli, The heterochromatin protein 1 prevents telomere fusions in Drosophila. Mol Cell 2, 527–538 (1998).

38. O. Komonyi, T. Schauer, G. Papai, P. Deak, I. M. Boros, A product of the bicistronic Drosophila melanogaster gene CG31241, which also encodes a trimethylguanosine synthase, plays a role in telomere protection. J Cell Sci 122, 769–774 (2009).

39. R. Dubruille et al., Specialization of a Drosophila capping protein essential for the protection of sperm telomeres. Curr Biol 20, 2090–2099 (2010).

40. G. Gao, Y. Cheng, N. Wesolowska, Y. S. Rong, Paternal imprint essential for the inheritance of telomere identity in Drosophila. Proc Natl Acad Sci U S A 108, 4932–4937 (2011).

41. T. Yamaki, G. K. Yasuda, B. T. Wakimoto, The Deadbeat paternal effect of uncapped sperm telomeres on cell cycle progression and chromosome behavior in Drosophila melanogaster. Genetics 203, 799–816 (2016).

42. B. Saint-Leandre, C. Christopher, M. T. Levine, Adaptive evolution of an essential telomere protein restricts telomeric retrotransposons. Elife 9, (2020).

43. K. B. S. Swamy, S. C. Schuyler, J. Y. Leu, Protein complexes form a basis for complex hybrid incompatibility. Front Genet 12, 609766 (2021).

44. S. De Renzis, O. Elemento, S. Tavazoie, E. F. Wieschaus, Unmasking activation of the zygotic genome using chromosomal deletions in the Drosophila embryo. PLoS Biol 5, e117 (2007).

45. S. J. Wright, Sperm nuclear activation during fertilization. Curr Top Dev Biol 46, 133–178 (1999).

46. B. Loppin, R. Dubruille, B. Horard, The intimate genetics of Drosophila fertilization. Open Biol 5, (2015).

47. G. W. van der Heijden et al., Asymmetry in histone H3 variants and lysine methylation between paternal and maternal chromatin of the early mouse zygote. Mech Dev 122, 1008–1022 (2005).

48. B. Loppin, D. Lepetit, S. Dorus, P. Couble, T. L. Karr, Origin and neofunctionalization of a Drosophila paternal effect gene essential for zygote viability. Curr Biol 15, 87–93 (2005).

49. M. Cui, Y. F. Bai, K. L. Li, Y. K. S. Rong, Taming active transposons at Drosophila telomeres: The interconnection between HipHop’s roles in capping and transcriptional silencing. Plos Genetics 17, (2021).

50. N. C. Elde, H. S. Malik, The evolutionary conundrum of pathogen mimicry. Nat Rev Microbiol 7, 787–797 (2009).

51. S. S. Parhad, S. Tu, Z. Weng, W. E. Theurkauf, Adaptive evolution leads to cross-species incompatibility in the piRNA transposon silencing machinery. Dev Cell 43, 60–70 e65 (2017).

52. H. A. Orr, Dobzhansky, Bateson, and the genetics of speciation. Genetics 144, 1331–1335 (1996).

53. D. C. Presgraves, C. D. Meiklejohn, Hybrid sterility, genetic conflict and complex speciation: Lessons from the Drosophila simulans clade species. Front Genet 12, 669045 (2021).

54. L. Chen et al., AI-driven deep learning techniques in protein structure prediction. Int J Mol Sci 25, (2024).

55. K. Tunyasuvunakool et al., Highly accurate protein structure prediction for the human proteome. Nature 596, 590–596 (2021).

56. G. Kim et al., Easy and accurate protein structure prediction using ColabFold. Nat Protoc, (2024).

57. E. L. van Dijk et al., Genomics in the long-read sequencing era. Trends Genet 39, 649–671 (2023).

58. H. Li, R. Durbin, Genome assembly in the telomere-to-telomere era. Nat Rev Genet 25, 658–670 (2024).

59. R. C. Edgar, Muscle5: High-accuracy alignment ensembles enable unbiased assessments of sequence homology and phylogeny. Nat Commun 13, 6968 (2022).

60. M. Suyama, D. Torrents, P. Bork, PAL2NAL: Robust conversion of protein sequence alignments into the corresponding codon alignments. Nucleic Acids Res 34, W609–W612 (2006).

61. Z. Yang, PAML 4: Phylogenetic analysis by maximum likelihood. Mol Biol Evol 24, 1586–1591 (2007).

62. J. H. Mcdonald, M. Kreitman, Adaptive protein evolution at the Adh Locus in Drosophila. Nature 351, 652–654 (1991).

63. J. B. Lack et al., The Drosophila genome nexus: A population genomic resource of 623 Drosophila melanogaster genomes, including 197 from a single ancestral range population. Genetics 199, 1229–1241 (2015).

64. J. Rozas et al., DnaSP 6: DNA sequence polymorphism analysis of large data sets. Mol Biol Evol 34, 3299–3302 (2017).

65. S. Kondo, R. Ueda, Highly improved gene targeting by germline-specific Cas9 expression in Drosophila. Genetics 195, 715–721 (2013).

66. S. J. Gratz et al., Highly specific and efficient CRISPR/Cas9-catalyzed homology-directed repair in Drosophila. Genetics 196, 961–971 (2014).

67. J. Gokcezade, G. Sienski, P. Duchek, Efficient CRISPR/Cas9 plasmids for rapid and versatile genome editing in Drosophila. G3 (Bethesda) 4, 2279–2282 (2014).

68. A. C. Groth, M. Fish, R. Nusse, M. P. Calos, Construction of transgenic Drosophila by using the site-specific integrase from phage phiC31. Genetics 166, 1775–1782 (2004).

69. W. F. Rothwell, W. Sullivan, Drosophila embryo collection. CSH Protoc 2007, pdb prot4825 (2007).

70. W. F. Rothwell, W. Sullivan, Fixation of Drosophila embryos. CSH Protoc 2007, pdb prot4827 (2007).

71. B. Loppin, F. Berger, P. Couble, The Drosophila maternal gene sesame is required for sperm chromatin remodeling at fertilization. Chromosoma 110, 430–440 (2001).

72. F. Ciabrelli, N. Atinbayeva, A. Pane, N. Iovino, Epigenetic inheritance and gene expression regulation in early Drosophila embryos. EMBO Rep 25, 4131–4152 (2024).

73. J. Schindelin et al., Fiji: An open-source platform for biological-image analysis. Nat Methods 9, 676–682 (2012).

