## Supplementary material for "Adaptive protein coevolution preserves telomere integrity": LinETAL_Supplement

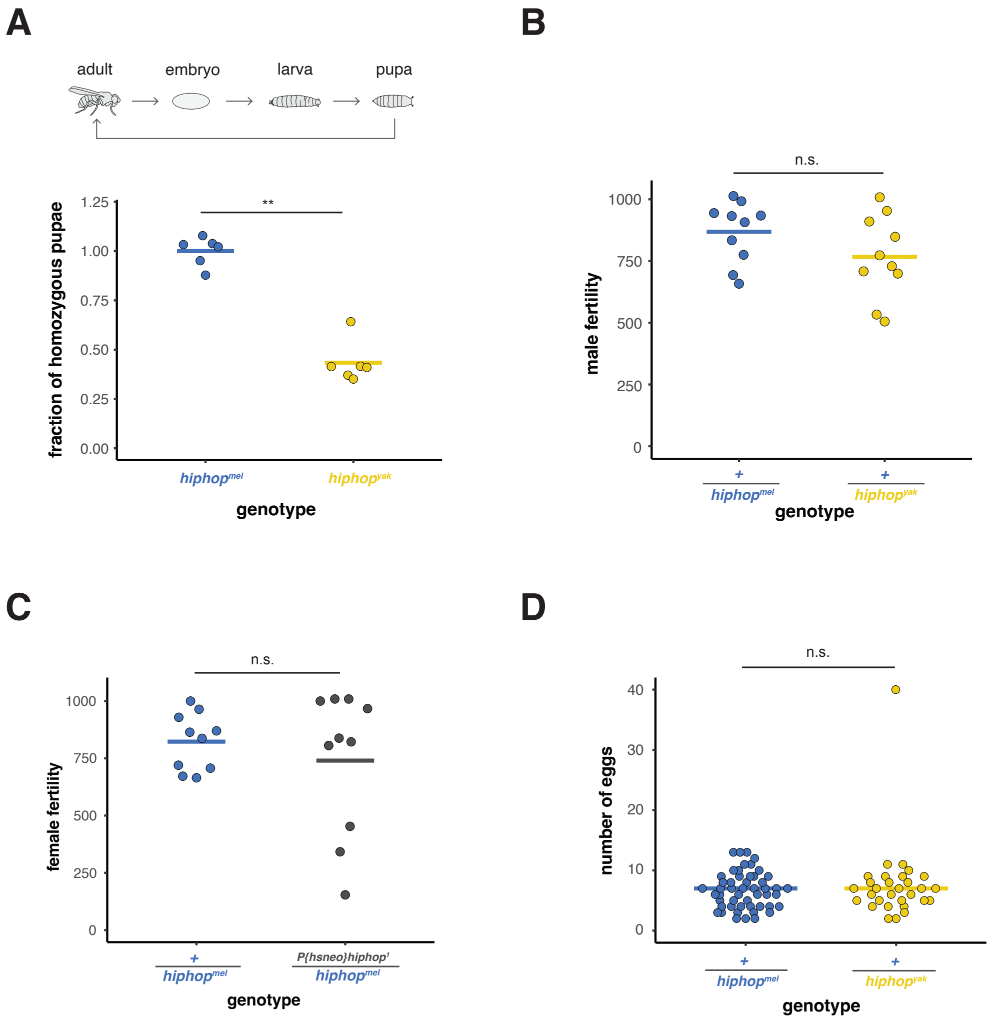


**Fig S1. HipHop$^{yak}$ causes pupal inviability and has no detectable impact on *D. melanogaster* male fertility or female fecundity.** (**A**) Schematic of the life cycle of *Drosophila* (above). Pupal viability is reduced in *hiphop$^{yak}$* homozygotes (below). Mann-Whitney U (MWU) test, ***p*-value < 0.01. (**B**) No difference in male fertility detected between the +/*hiphop$^{mel}$* and +/*hiphop$^{yak}$* genotypes. "+" = *TM6* balancer chromosome. MWU test. (**C**) No reduction in female fertility detected in the *hiphop* null mutant. *P{hsneo}hiphop$^{1}$* = *hiphop* null mutant (*1*). MWU test. (**D**) No difference in female fecundity detected between the +/*hiphop$^{mel}$* and +/*hiphop$^{yak}$* genotypes. "+" = *TM6* balancer chromosome. n ≥ 30. MWU test.

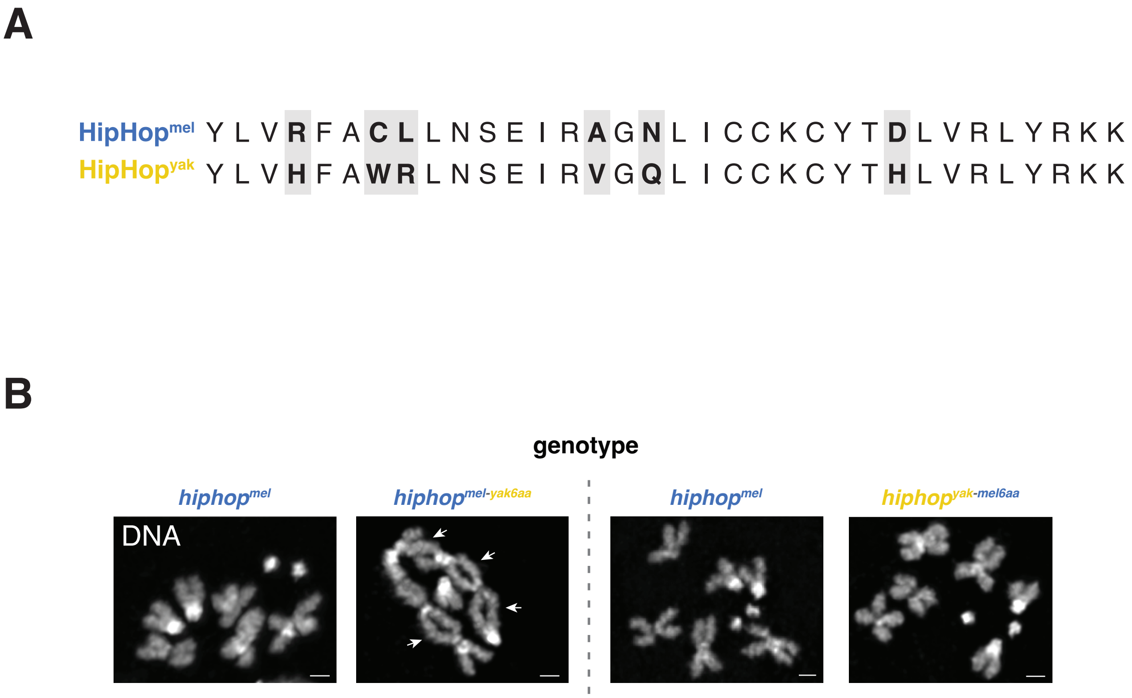


**Fig S2. Evolution of at most six amino acids along the HipHop-HOAP interaction surface is necessary and sufficient for chromosome end-protection.** (**A**) Amino acid alignment of the positively selected region between HipHop$^{mel}$ and HipHop$^{yak}$. The shaded boxes indicate the six diverged residues in Fig. 2D and 4A. (**B**) Mitotic chromosomes from larval brains stained with DAPI demonstrate that evolution of at most six residues on the HipHop-HOAP interaction surface preserves end-protection. n ≥ 58 mitotic cells.

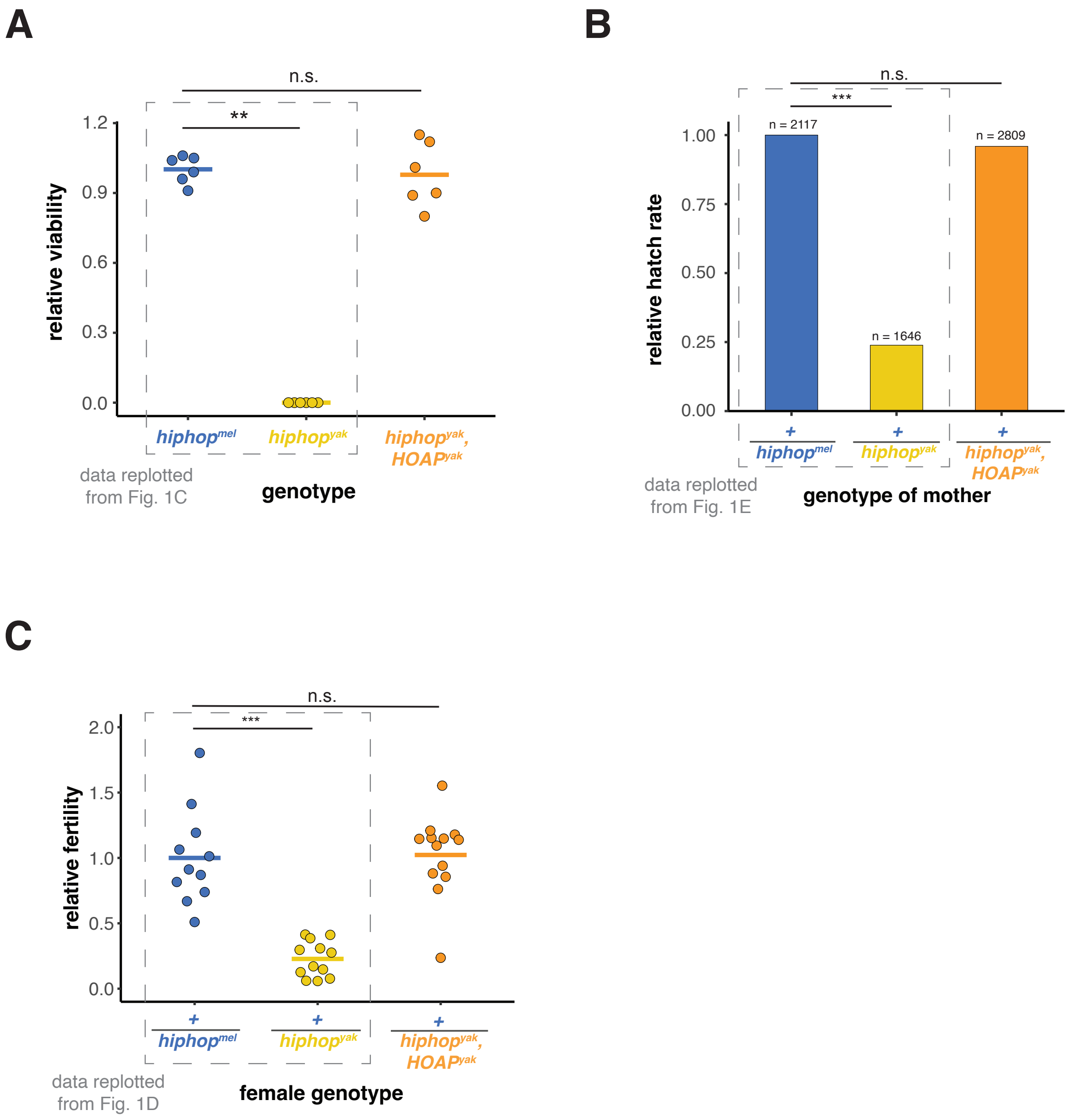


**Fig S3. *HOAP*$^{yak}$ rescues viability, embryo hatch rate, and female fertility of *hiphop*$^{yak}$. (A)** Relative viability of *hiphop*$^{mel}$, *hiphop*$^{yak}$, and *hiphop*$^{yak}$*,HOAP*$^{yak}$ (*hiphop*$^{mel}$ and *hiphop*$^{yak}$ data replotted from Fig. 1C). Mann-Whitney U test: ***p*-value < 0.01. (**B**) Relative hatch rate of embryos from +/*hiphop*$^{mel}$, +/*hiphop*$^{yak}$, and +/*hiphop*$^{yak}$*,HOAP*$^{yak}$ females (+/*hiphop*$^{mel}$ and +/*hiphop*$^{yak}$ data replotted from Fig. 1E). Fisher's Exact Test: ****p*-value < 0.001. "+" = *TM6* balancer chromosome. (**C**) The relative female fertility of +/*hiphop*$^{mel}$, +/*hiphop*$^{yak}$, and +/*hiphop*$^{yak}$*,HOAP*$^{yak}$ females crossed to wild-type (*w1118*) males (+/*hiphop*$^{mel}$ and +/*hiphop*$^{yak}$ data replotted from Fig. 1D). MWU: ****p*-value < 0.001.

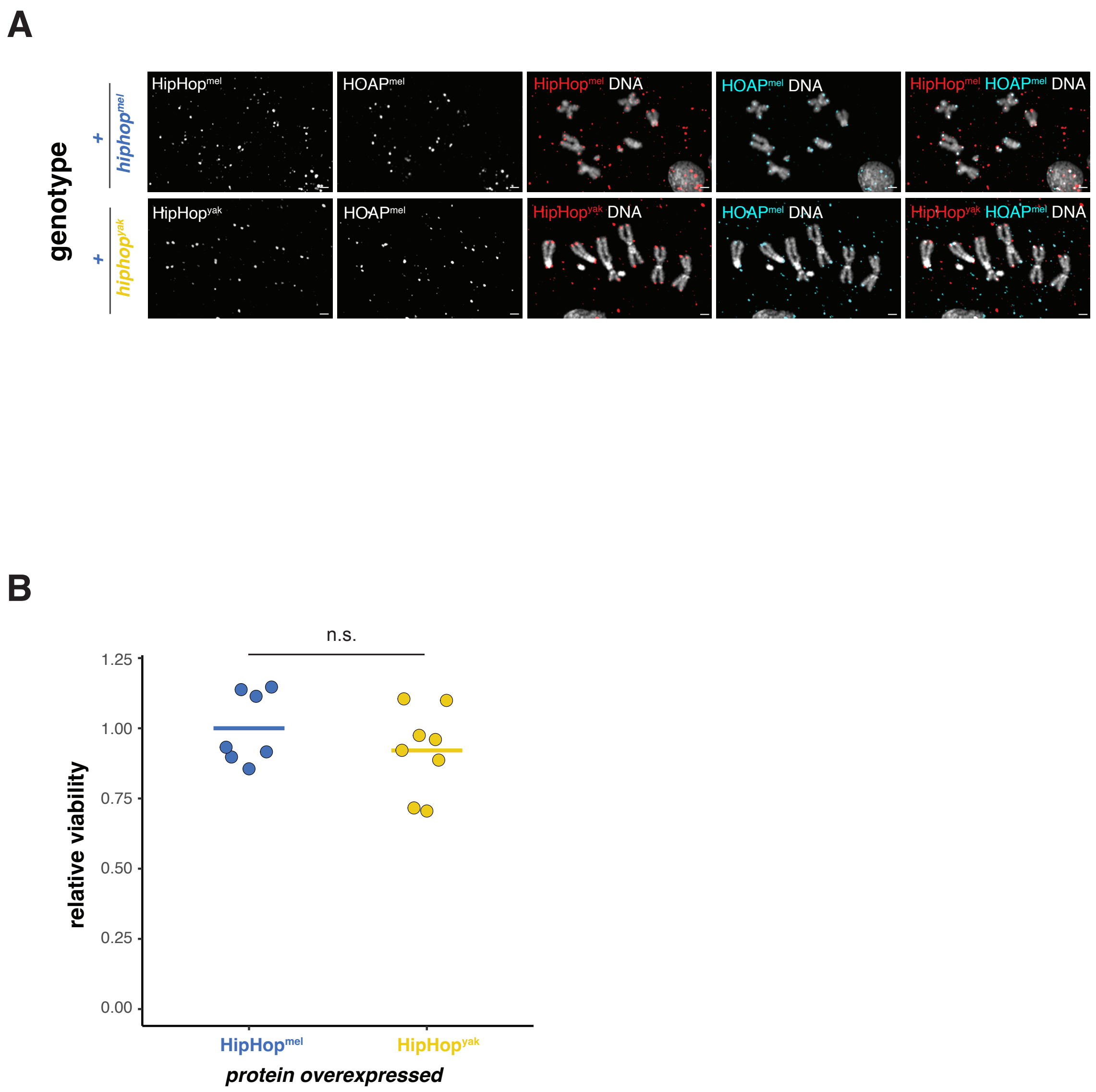


**Fig S4. Once end-protection is established, telomere integrity and viability are preserved even in the presence of HipHop$^{yak}$.** (**A**) Chromosome end-protection and telomere localization of HOAP are maintained in the +/*hiphop$^{yak}$* larvae. The well-resolved, mitotic chromosomes from larval brains stained for HipHop (anti-EGFP) and HOAP (anti-FLAG) in +/*hiphop$^{mel}$* and +/*hiphop$^{yak}$* heterozygotes. "+" = *TM6* balancer chromosome. (**B**) No reduction in relative viability detected upon ubiquitous overexpression of HipHop$^{yak}$. HipHop$^{mel}$ = *UAS-hiphop$^{mel}$/+; Act5C-GAL4/+* and HipHop$^{yak}$ = *UAS-hiphop$^{yak}$/+; Act5C-GAL4/+*. MWU test.

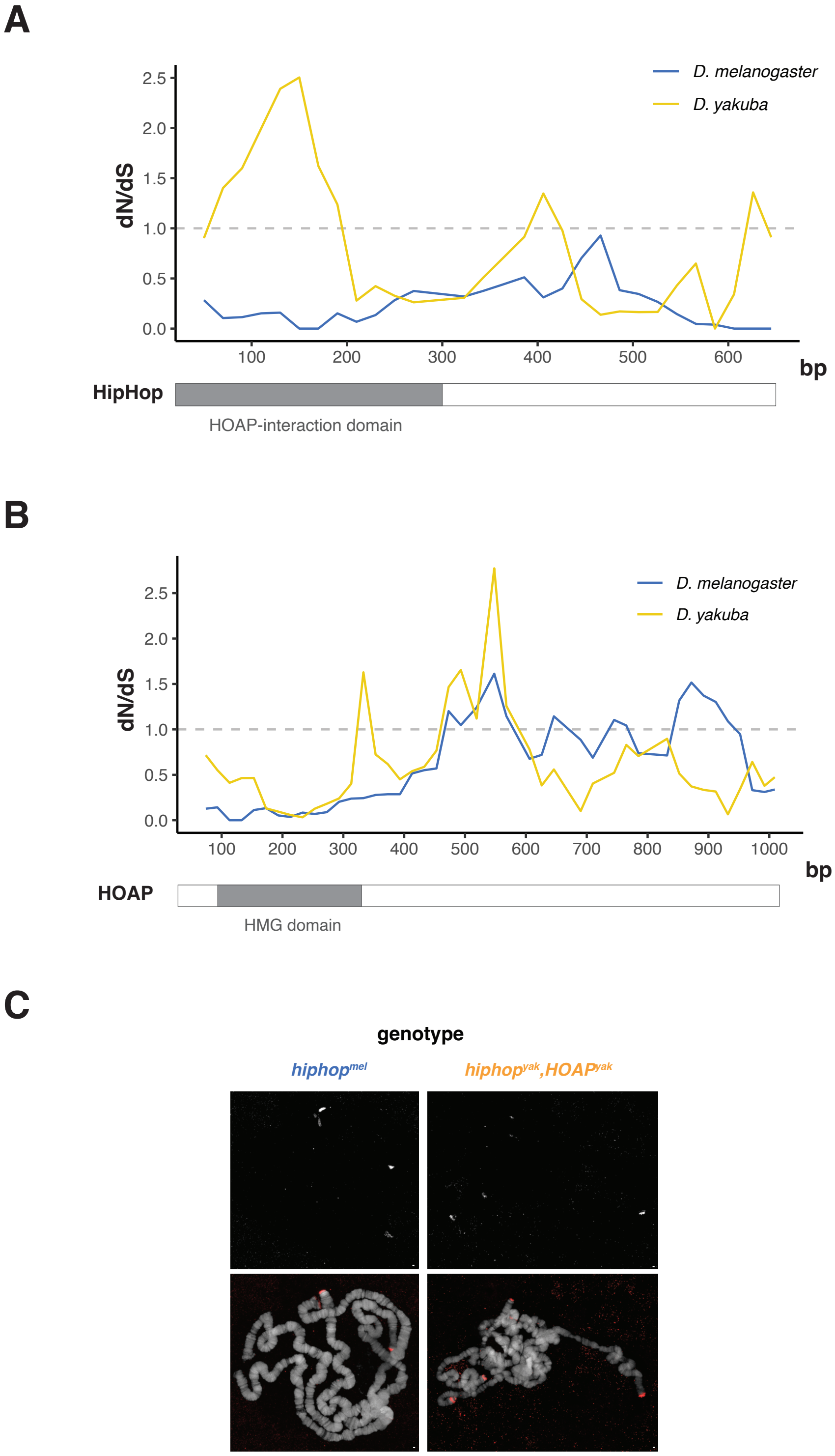
A
D. melanogaster
D. yakuba
dN/dS
2.5
2.0
1.5
1.0
0.5
0.0
100
200
300
400
500
600
bp
HipHop
HOAP-interaction domain
B
D. melanogaster
D. yakuba
dN/dS
2.5
2.0
1.5
1.0
0.5
0.0
100
200
300
400
500
600
700
800
900
1000
bp
HOAP
HMG domain
C
genotype
hiphop^mel
hiphop^yak,HOAP^yak

**Fig S5. Support for the proposed model of retrotransposon-triggered protein-protein coevolution.** (**A** and **B)** The estimated rate of lineage-specific nonsynonymous substitutions relative to synonymous substitutions (dN/dS) across the *D. melanogaster* (blue) and *D. yakuba* (yellow) *hiphop* (A) and *cav/HOAP* (B) coding sequences (window size = 100, step size = 20). The HMG domain mediates DNA-binding. (**C**) No evidence of telomere elongation in the *hiphop$^{yak}$,HOAP$^{yak}$* strain that evolved in the lab for 30 generations. FISH probe cognate to the HeT-A telomeric retrotransposon hybridized to polytene chromosomes from *hiphop$^{mel}$* or *hiphop$^{yak}$* flies.

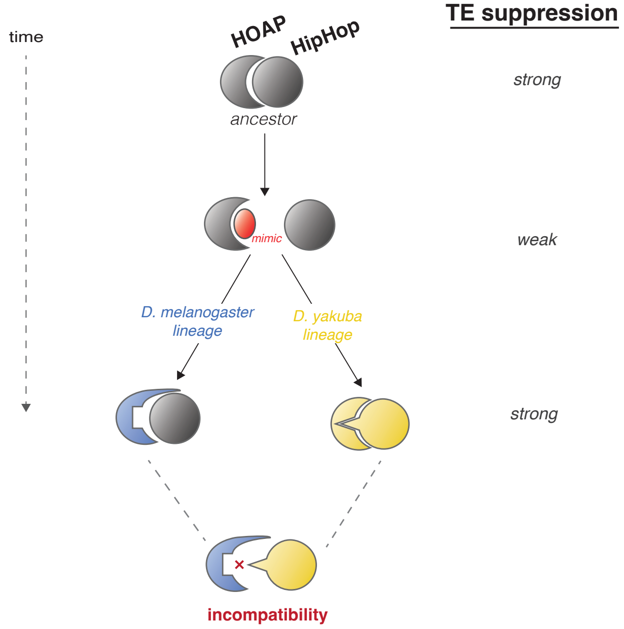


**Fig S6. A model of protein-protein coevolution triggered by retrotransposons.** We propose that fast-evolving telomeric retrotransposons trigger HipHop-HOAP coevolution. Under this model, telomeric retrotransposons evolve to escape host restriction by encoding a mimic (*3, 4*) of the HipHop-HOAP interaction surface. This mimic reduces the abundance of the HipHop-HOAP-HP1 subcomplex at telomeres, the dose of which may be critical for restricting telomere-specialized retrotransposons (*2, 5*). A fitness reduction selects for HOAP substitutions at the interaction surface along both lineages to avoid recognition. Along the *D. yakuba* lineage, HOAP evolution compromises its interaction with HipHop. Reduced interaction triggers HipHop lineage-specific evolution at HipHop's HOAP-interaction surface to preserve the abundance of this subcomplex. Along the *D. melanogaster* lineage, we detected no evidence of HipHop adaptive evolution. This distinct evolutionary trajectory along the *D. melanogaster* lineage suggests that HOAP evolved at multiple surfaces to avoid the mimic but preserve its interaction with HipHop. Previous work found that swapping *HOAP$^{yak}$* into *D. melanogaster* does not cause lethality but does activate transposons (*2*), raising the possibility that along both species' lineages, retrotransposon restriction is restored by the adaptive evolution of one or both proteins. Indeed, we observe no evidence of telomere elongation in the *hiphop$^{yak}$,HOAP$^{yak}$* genotype (Fig. S5C), suggesting that *hiphop$^{yak}$* rescues telomeric retrotransposon restriction, possibly by restoring the abundance of the HipHop-HOAP-HP1 subcomplex. This model accounts not only for the distinct lineage-specific molecular evolution of these two proteins (see S5A and S5B) but also for the non-reciprocal lethal incompatibility of swapping the *D. yakuba* versions of *hiphop* and *HOAP* (*2*). Swapping *hiphop$^{yak}$* into *D. melanogaster* is lethal but swapping *HOAP$^{yak}$* into *D. melanogaster* is not. Future work will probe the mechanism by which telomeric retrotransposons antagonize this protein-protein interaction.

**Table S1. HipHop and HOAP evolve adaptively in the *melanogaster* subgroup.** A phylogeny-based test using the codeml program in PAML was performed on genes that make up the *Drosophila* chromosome end-protection complex. The analysis included only the *melanogaster* subgroup (*D. melanogaster*, *D. simulans*, *D. sechellia*, *D. mauritiana*, *D. erecta*, *D. yakuba*, and *D. teissieri*).

| gene | # codon analyzed | M7 lnL | M8 lnL | log likelihood ratio | *p* value | positively selected sites (probability > 95%) |
|---|---|---|---|---|---|---|
| ***hiphop*** | ***219*** | ***-2003.08*** | ***-1997.06*** | ***12.04*** | ***0.002*** | ***154L**** |
| ***cav/HOAP*** | ***309*** | ***-2640.61*** | ***-2636.86*** | ***7.50*** | ***0.024*** | ***159S*, 160N**** |
| *Su(var)205* | 205 | -1318.69 | -1318.69 | 0.00 | 1.000 | NA |
| *ver* | 214 | -1780.56 | -1779.96 | 1.20 | 0.549 | NA |
| *moi* | 166 | -1114.95 | -1115.00 | 0.10 | 0.949 | NA |
| *tea* | 1792 | -15732.83 | -15731.56 | 2.56 | 0.278 | NA |

**Table S2. HipHop evolves adaptively between *D. melanogaster* and *D. yakuba*.** Counts of synonymous and nonsynonymous polymorphic and diverged sites of *hiphop* coding region within and between *D. melanogaster* and *D. yakuba*. Fisher's Exact test, $p = 0.06$.

| | polymorphism | divergence |
|---|---|---|
| synonymous | 14 | 51 |
| nonsynonymous | 7 | 65 |

### Literature Cited


1. R. Dubruille *et al.*, Specialization of a *Drosophila* capping protein essential for the protection of sperm telomeres. *Curr Biol* **20**, 2090-2099 (2010).
2. B. Saint-Leandre, C. Christopher, M. T. Levine, Adaptive evolution of an essential telomere protein restricts telomeric retrotransposons. *Elife* **9**, (2020).
3. S. S. Parhad, S. Tu, Z. Weng, W. E. Theurkauf, Adaptive evolution leads to cross-species incompatibility in the piRNA transposon silencing machinery. *Dev Cell* **43**, 60-70 e65 (2017).
4. N. C. Elde, H. S. Malik, The evolutionary conundrum of pathogen mimicry. *Nat Rev Microbiol* **7**, 787-797 (2009).
5. M. Cui, Y. F. Bai, K. L. Li, Y. K. S. Rong, Taming active transposons at *Drosophila* telomeres: The interconnection between HipHop's roles in capping and transcriptional silencing. *Plos Genetics* **17**, (2021).
